# The nuclear receptor LRH-1 discriminates between ligands using distinct allosteric signaling circuits

**DOI:** 10.1101/2023.01.27.525934

**Authors:** Suzanne G. Mays, David Hercules, Eric A. Ortlund, C. Denise Okafor

**Author notes:** Department of Genome Biology, Centre for Genomic Regulation, Calle Dr. Aiguader 88, Barcelona, 08003, Spain.

## Abstract

Nuclear receptors (NRs) are transcription factors that regulate essential biological processes in response to cognate ligands. An important part of NR function involves ligand-induced conformational changes that recruit coregulator proteins to the activation function surface (AFS), ~15 Å away from the ligand binding pocket. Ligands must communicate with the AFS to recruit appropriate coregulators and elicit different transcriptional outcomes, but this communication is poorly understood. These studies illuminate allosteric communication networks underlying activation of liver receptor homolog-1 (LRH-1), a NR that regulates development, metabolism, cancer progression and intestinal inflammation. Using >100 microseconds of all-atom molecular dynamics simulations involving 69 LRH-1 complexes, we identify distinct signaling circuits used by active and inactive ligands for AFS communication. Inactive ligands communicate via strong, coordinated motions along paths through the receptor to the AFS. Activating ligands disrupt the “inactive” circuit by inducing connectivity elsewhere. Ligand-contacting residues in helix 7 help mediate the switch between circuits, suggesting new avenues for developing LRH-1-targeted therapeutics. We also elucidate aspects of coregulator signaling, showing that localized, destabilizing fluctuations are induced by inappropriate ligand-coregulator pairings. These studies have uncovered novel features of LRH-1 allostery, and the quantitative approach used to analyze many simulations provides a framework to study allosteric signaling in other receptors.

## Introduction

Nuclear receptors (NRs) are ligand-regulated transcription factors that control critical biological processes such as development, growth, and metabolism. Mechanisms of NR activation are complex, often involving simultaneous activation and repression of target genes in response to ligand binding. In addition, NR activation is modulated by coregulator proteins, which bind NR and regulate recruitment of the general transcriptional machinery. Therefore, NRs rely upon coordination of ligand- and coregulator-derived signals to execute coherent transcriptional programs.

NRs are excellent drug targets due to the presence of a defined ligand binding pocket and the ability to modulate gene programs that are often tissue-specific and linked to many diseases (1–5). Some NRs, such as the glucocorticoid receptor and estrogen receptor, have been targeted extensively with small molecules (6–8). The therapeutic potential of many other NRs remains largely untapped, in part due to an incomplete understanding of mechanisms of their regulation by ligands. This is the case for liver receptor homolog-1 (LRH-1), a therapeutic target for nonalcoholic fatty liver disease, diabetes, cancer, and inflammatory bowel diseases (9–12). LRH-1 regulates processes such as nutrient usage and gut inflammation by directing lipid, bile acid, and methyl metabolism in the liver (11,13–16), glucose uptake in skeletal muscle (17), and steroid biosynthesis in the intestine (9,18). Understanding mechanisms through which ligands switch LRH-1 into a transcriptionally active state could inform the design of LRH-1 modulators and also offer more general insights into NR regulation by ligands.

NR activation or inhibition involves allosteric communication between the ligand and the activation function surface (AFS), the site of coregulator binding (Figure 1A). For many NRs, this communication results in a dramatic conformational change to a helix in the AFS termed the activation function helix (AF-H) (19). LRH-1 is unconventional: its activation is thought to involve subtle, rather than large, conformational changes to the AFS, coupled with fluctuations at a second regulatory surface known as AFB (comprised of helix 6 and two beta strands, Figure 1A) (20). Loops flanking AFB impart flexibility to the surface, and this flexibility is required for LRH-1 activation by endogenous ligands (20). Further supporting the role of AFB in LRH-1 activation, strong communication occurs between AFB and the AFS in molecular dynamics (MD) simulations when an agonist is bound in the presence of an activating coregulator (“coactivator”) (21). Likewise, apo-LRH-1 paired with an inactivating coregulator (“corepressor”) strengthens communication between the two regulatory surfaces. However, when the ligand and coregulator are “mismatched” (*e.g*., with an agonist and a corepressor), the communication is weakened (21). While compelling, the studies identifying this communication only involved a handful of agonists and did not include inactive ligands, relying on apo-LRH-1 to model the inactive state.

**Figure 1.**
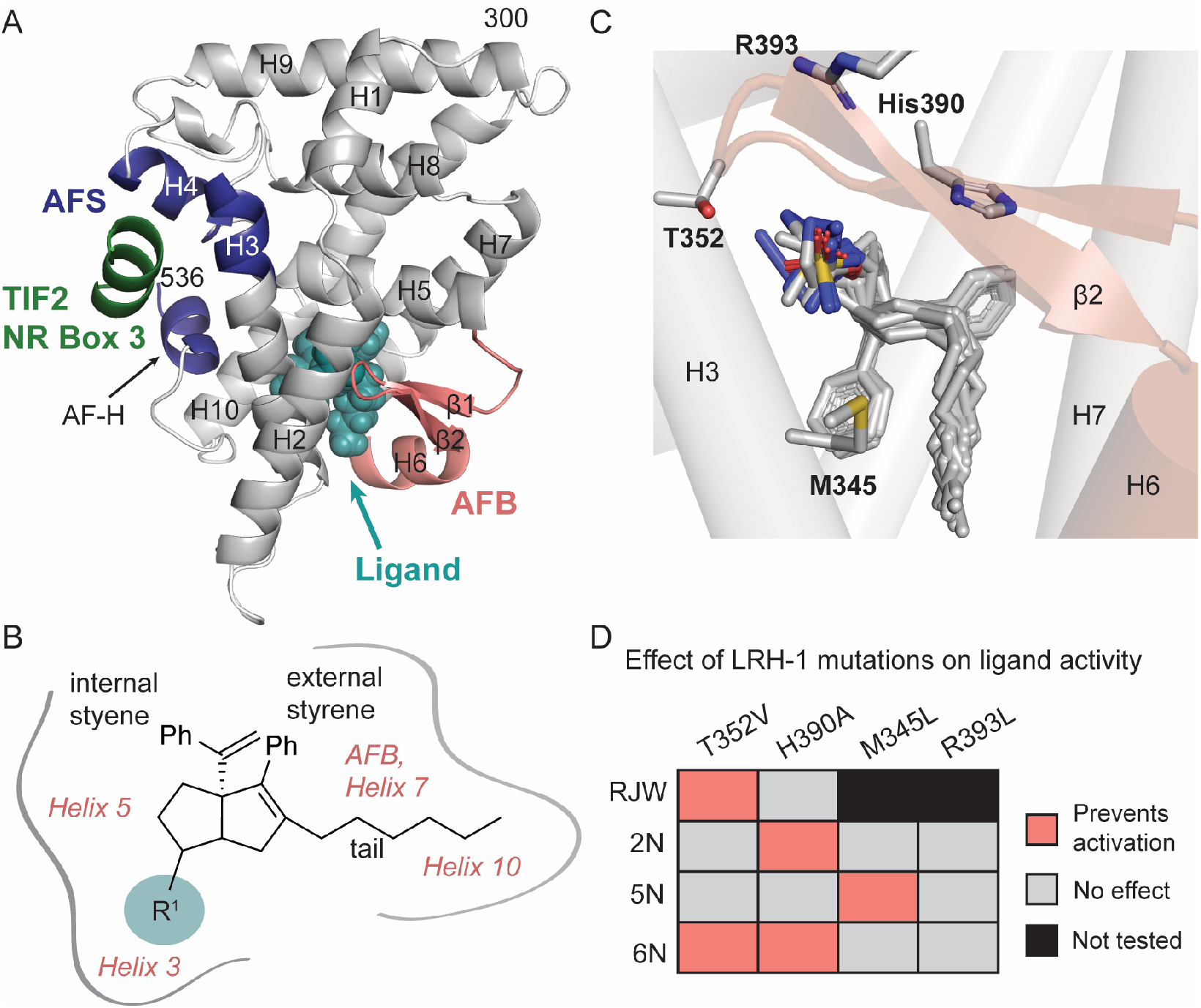
LRH-1 ligand binding domain and interactions with synthetic ligands. A. Model of the LRH-1 ligand binding domain (from PDB 6OQY), highlighting regions of interest. A ligand is bound in the ligand binding pocket (represented by teal spheres). The two allosteric surfaces discussed in this paper are also highlighted: the AFS is colored dark blue, and AFB is salmon. The Tif2 peptide containing the NR Box 3 interaction motif is shown bound at the AFS (dark green helix). B.6HP chemical scaffold of the ligands used in the MD simulations. The modified R^1^ site is highlighted in teal (modifications are shown in Table 1) Features common to all ligands are labeled in black, and the regions of LRH-1 making contact with the ligands are indicated in salmon. C. Close view of the LRH-1 ligand binding pocket, showing a superposition of the ligands used in the simulations. Residues previously shown to be important for activation for some of the ligands are shown as sticks (labeled in bold type). Helices are shown as cylinders, abbreviated “H”. AFB is colored pale salmon. D. Heatmap depicting the effect of the indicated point mutations on LRH-1 activation by ligands in this study.

In these studies, we probe mechanisms of LRH-1 activation with MD simulations involving 24 ligands: 12 that activate LRH-1 and 12 that bind LRH-1 but are inactive in luciferase reporter assays. The ligands were selected from a library of compounds developed in our laboratory containing the hexahydropentelene (6HP) scaffold and modified at a single site (position R^1^, Figure 1B)(22). The modifications were aimed at introducing interactions with LRH-1 residues in a polar patch of the binding pocket near helices 3 and 5 (Figure 1C). The 6HP ligands selected for MD simulations demonstrate a strong structure-activity relationship: certain polar groups at R^1^ maintain or increase the activity of the parent compound and increase potency by up to two orders of magnitude (22,23). Structural and functional studies have identified interactions required for LRH-1 activation, which differ slightly for different ligands (Figure 1D). The availability of robust biological and mechanistic data for these compounds makes them ideal to study dynamics underlying ligand activation.

An analysis of LRH-1 conformational dynamics in the simulations reveals a central role for the N-terminal part of helix 7 in communicating ligand status to the AFS. Inactive ligands promote communication between helix 7 and the AFS, while active ligands disrupt helix 7-AFS communication by inducing connectivity between the ligand, helix 7 and AFB. The addition of a coactivator to complexes containing inactive ligands causes fluctuations within the ligands and portions of the receptor, which are strongest in AFB. We propose that inactive ligands send a “stop” signal to the AFS to disfavor coactivator binding, and that this signal is suppressed in the presence of active ligands. Forcing a coactivator to bind with an inactive ligand causes disharmony in the receptor that is reflected in AFB fluctuations.

## Results

### Ligand classification

To uncover molecular motions driving LRH-1 activation, we conducted MD simulations with 69 different LRH-1 complexes involving 24 ligands (Table S1). The ligands used in the simulations are closely related but have diverse biological profiles, ranging from completely inactive to highly active. In general, incorporation of tetrahedral polar groups at position R^1^ in the 6HP scaffold (Figure 1B) improves binding affinity, increases LRH-1 thermostability by differential scanning fluorimetry (DSF), and increases LRH-1 activity in luciferase reporter assays (biological data and ligand chemical structures are summarized in Table 1) (22). We considered both potency (EC_50_) and efficacy (E_max_) from luciferase reporter assays measuring LRH-1 activity to classify the ligands into two main groups. The “active” group (n = 12) contains ligands that dose-dependently activate LRH-1, produce well-defined plateaus at the tops of dose-response curves, and have EC_50_s in the nanomolar to low micromolar range(22). The “inactive” group contains 12 ligands that bind LRH-1 but do not activate it at the highest dose tested (30 μM) or for which an EC_50_ could not be calculated due to low potency. We constructed models for MD simulations using published crystal structures of LRH-1 bound to 6HP ligands (PBD 5L11, 5SZY, 6OR1, 6OQX, and 6OQY)(22,24). We have shown that polar groups at R1 in 6HP ligands act as anchors to lock this class of compound in very similar binding poses. (22,24–27). Therefore, ligands without experimentally determined structures were docked into the binding pocket such that the 6HP cores were aligned (Figure 1C). We also evaluated LRH-1 with no ligand bound (apo-LRH-1), using PDB 4PLD as the starting model (21). All structures selected as models were crystallized with a fragment of the coactivator protein transcriptional intermediary factor 2 (Tif2) bound at the LRH-1 AFS, and we conducted simulations with and without Tif2 to investigate how coregulator binding affects LRH-1 conformational dynamics in the presence of different ligands. For each complex, 500 ns of simulations were conducted in triplicate, and data from the triplicate runs were averaged. Previous studies show that compared to one long continuous trajectory, replicas of shorter MD simulations reduce the likelihood of false positive conclusions (28).

**Table 1.**
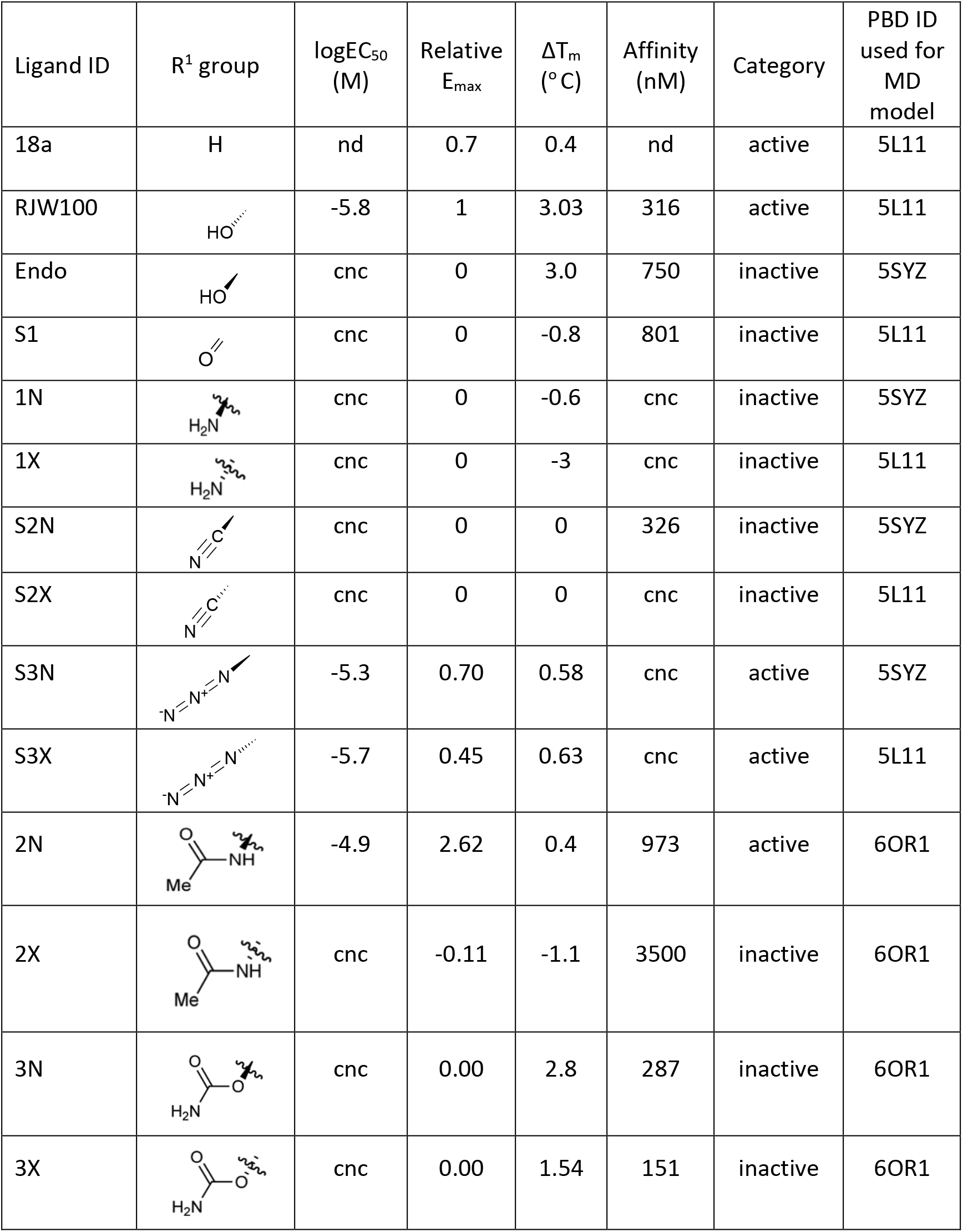
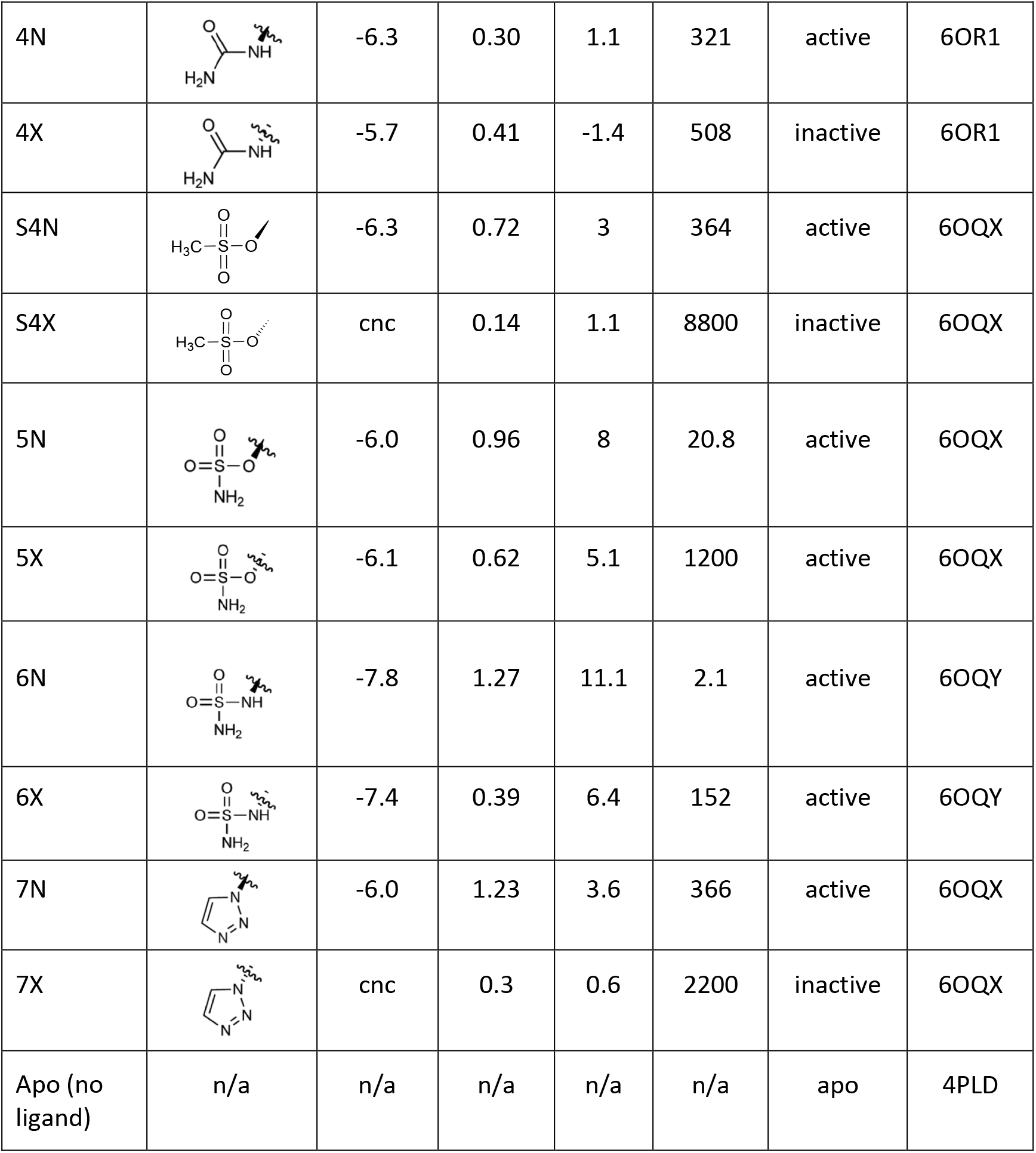
Compounds used for molecular dynamics (MD) simulations and biological profiles from previous studies (22,24,29). cnc, could not calculate; nd, not done; n/a, not applicable. For a list of all simulations run (including +/- coactivator and for LRH-1 mutants) see Table S1.

| Ligand ID | R<sup>1</sup> group | logEC<sub>50</sub><br>(M) | Relative<br>E<sub>max</sub> | ΔT<sub>m</sub><br>(° C) | Affinity<br>(nM) | Category | PBD ID<br>used for<br>MD<br>model |
| --- | --- | --- | --- | --- | --- | --- | --- |
| 18a | H | nd | 0.7 | 0.4 | nd | active | 5L11 |
| RJW100 |  | -5.8 | 1 | 3.03 | 316 | active | 5L11 |
| Endo |  | cnc | 0 | 3.0 | 750 | inactive | 5SYZ |
| S1 |  | cnc | 0 | -0.8 | 801 | inactive | 5L11 |
| 1N |  | cnc | 0 | -0.6 | cnc | inactive | 5SYZ |
| 1X |  | cnc | 0 | -3 | cnc | inactive | 5L11 |
| S2N |  | cnc | 0 | 0 | 326 | inactive | 5SYZ |
| S2X |  | cnc | 0 | 0 | cnc | inactive | 5L11 |
| S3N |  | -5.3 | 0.70 | 0.58 | cnc | active | 5SYZ |
| S3X |  | -5.7 | 0.45 | 0.63 | cnc | active | 5L11 |
| 2N |  | -4.9 | 2.62 | 0.4 | 973 | active | 6OR1 |
| 2X |  | cnc | -0.11 | -1.1 | 3500 | inactive | 6OR1 |
| 3N |  | cnc | 0.00 | 2.8 | 287 | inactive | 6OR1 |
| 3X |  | cnc | 0.00 | 1.54 | 151 | inactive | 6OR1 |
| 4N |  | -6.3 | 0.30 | 1.1 | 321 | active | 6OR1 |
| 4X |  | -5.7 | 0.41 | -1.4 | 508 | inactive | 6OR1 |
| S4N |  | -6.3 | 0.72 | 3 | 364 | active | 6OQX |
| S4X |  | cnc | 0.14 | 1.1 | 8800 | inactive | 6OQX |
| 5N |  | -6.0 | 0.96 | 8 | 20.8 | active | 6OQX |
| 5X |  | -6.1 | 0.62 | 5.1 | 1200 | active | 6OQX |
| 6N |  | -7.8 | 1.27 | 11.1 | 2.1 | active | 6OQY |
| 6X |  | -7.4 | 0.39 | 6.4 | 152 | active | 6OQY |
| 7N |  | -6.0 | 1.23 | 3.6 | 366 | active | 6OQX |
| 7X |  | cnc | 0.3 | 0.6 | 2200 | inactive | 6OQX |
| Apo (no ligand) | n/a | n/a | n/a | n/a | n/a | apo | 4PLD |

### Activating ligands form productive interactions with LRH-1 helix 7

To discover differences between active and inactive LRH-1 ligands, we first investigated receptor-ligand interactions and local conformational perturbations near the binding pocket. We focused on ligand-induced effects by analyzing complexes without a coregulator bound. There are no obvious differences in the stability of ligand interactions or which residues are involved: ligands in both classes interact with residues in helices 3, 5, 6, and 7 for greater than 90% of the simulation time (Figure S1A). Any differences in interaction stability appear to be related more to R^1^ group size than activity, since ligands with large R^1^ groups cluster together whether they are active or inactive (Figure S1A). However, active and inactive ligands have different effects on motions of residues they contact. We determined this by measuring coupled motion between ligands and each interacting residue, reasoning that close coupling likely reflects productive interactions related to receptor function. We quantified coupled motion using the value ‘shortest distance,’ which is inversely proportional to the correlation coefficient *r* between the ligand and the Cα of the interacting residue. Unbiased hierarchical clustering of the shortest distance data results in three main clusters containing ligands with distinct biological profiles (Figure 2A). Cluster 2 is particularly interesting: it contains five of the most potent active ligands, and it excludes inactive ligands with large R^1^ groups, supporting the idea that this analysis captures functional interactions (Figure 2A, inset). Ligands in cluster 2 also strongly stabilize LRH-1 in DSF experiments (measured by ΔTm), which was linked to potency in a previous study (22). Cluster 3 contains a mixture of active and inactive ligands that are less stabilizing by DSF than those in cluster 2, while cluster 1 contains mainly inactive ligands with smaller R^1^ groups (Figure 2A). The clustering is mainly driven by strong receptorligand coupling with residues in helix 3 (clusters 2 and 3) or with residues in helices 6 and 7 in addition to helix 3 (cluster 2, red box). Additional analyses of the shortest distance data for the full set of ligands reveal that potency (EC_50_) is correlated with stronger coupling of ligands with helix 3 (Figure 2B), consistent with experimental data linking potency to helix 3 interactions (22). Coupled motion between ligands and residues in helices 6, 7, and 10 is correlated with higher levels of activity in luciferase reporter assays (E_max_, Figure 2B).

**Figure 2.**
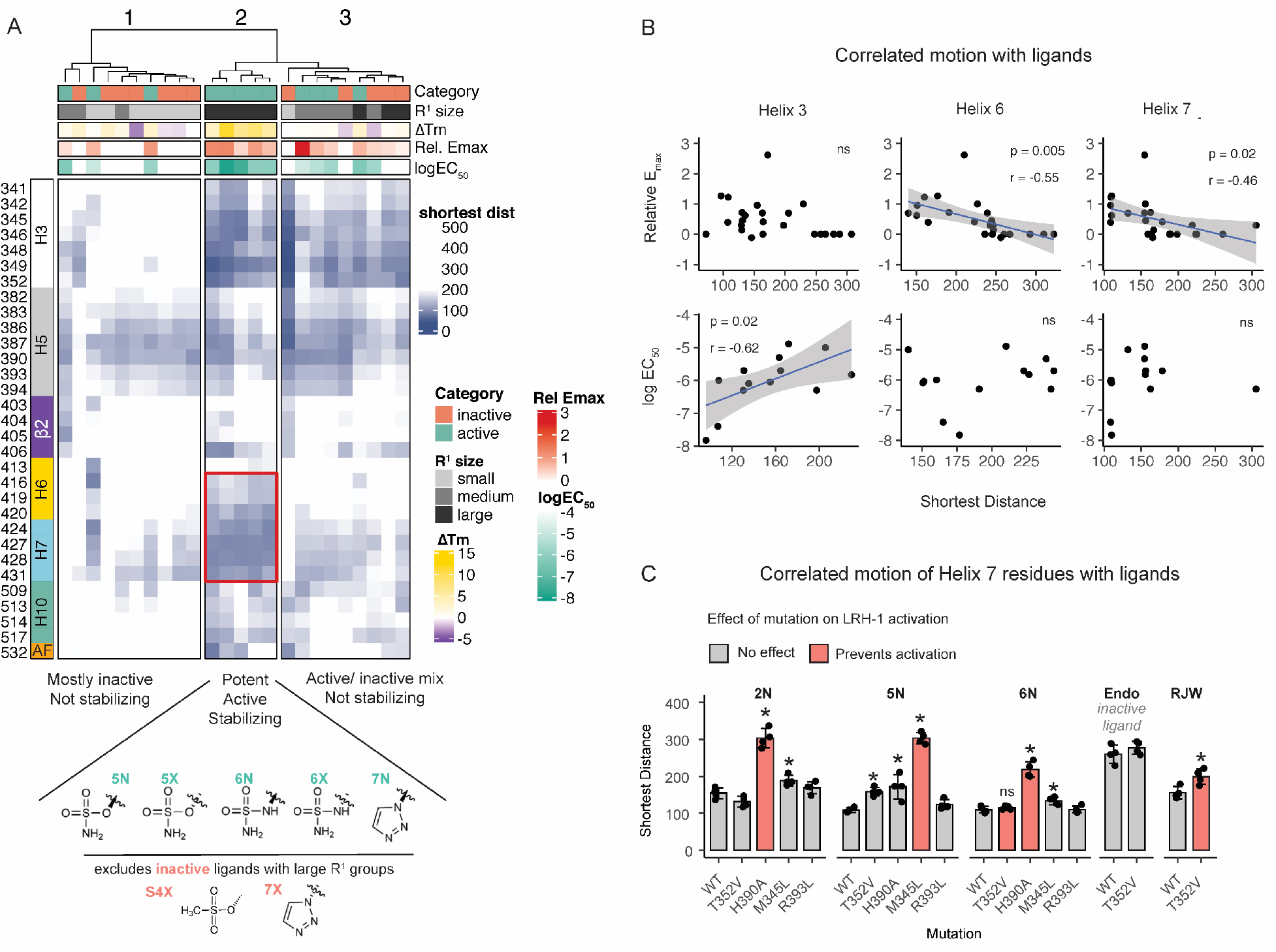
Activating ligands move with LRH-1 helix 7. A. Heatmap quantifying correlated motion between each ligand and the LRH-1 residues that they contact, measured by the value ‘shortest distance.’ Each column represents a ligand, and each row is an amino acid (numbered on the far left of the heatmap). Lower values for shortest distance (darker color) indicate stronger correlation between motions of the ligand and motions of the residue. Data are clustered by columns, using Euclidean distance and Ward’s agglomeration. Top annotation describes physical and biological characteristics of the ligands, including (1) relative E_max_ (maximum transcriptional activity in luciferase reporter assays relative to the parent compound, RJW100); (2) logEC_50_ from luciferase reporter assays, colored white if values could not be calculated due to poor activity); (3) ΔT_m_ (change in melting temperature for the protein-ligand complex upon ligand binding); (4) size of the R1 group, and (5) ligand classification used for this study. The color bar to the left of the heatmap describes the location of the LRH-1 residues (H, helix; β, beta-sheet). *Inset*, R^1^ groups of the ligands in cluster 2, as well as ligands with large R^1^ groups that are excluded from the cluster). This highlights the fact that the shortest distance analysis discriminates between active and inactive ligands with large R^1^ groups. B. Scatter plots describing the relationship between shortest distance value for residues in each helix (mean) and biological data for all ligands. For logEC_50_ graphs, ligands for which EC_50_s could not be calculated are omitted. The best-fit line (blue) and 95% CI (grey shading) are shown for linear relationships that were significant by Pearson’s correlation analysis. *Insets*, p-values and Pearson’s correlation coefficients (*r*) for correlations with p < 0.05; ns indicates not significant. C. Comparisons of shortest distance values (mean) for residues in helix 7 for wild-type LRH-1 (WT) or LRH-1 with the indicated point mutations. *, p < 0.05 relative to WT for each ligand (one-way ANOVA, Dunnett’s multiple comparisons test).

Helices 6 and 10 are known to be important for LRH-1 activity, but the involvement of helix 7 has not been described. We used *in silico* mutagenesis to further explore how its motion is linked to LRH-1 activation. We have identified mutations that prevent LRH-1 activation by particular ligands (summarized in Figure 1D) and investigated whether they could disrupt ligand-helix 7 coupling. Marked decoupling is seen for four out of five inactivating mutant-ligand combinations (Figure 2C, red bars). As a control, we also investigated ligand-mutation combinations that are not inactivating (Figure 2C, grey bars). In a few cases, ligand-helix 7 decoupling occurs to a level of statistical significance; however, the magnitude of the effect is smaller. The inactivating mutations have lesser effects or less specific effects on ligand coupling with other helices contacting the ligands (Figure S1B). Therefore, even though all the ligands interact stably with the same residues in helix 7 (Figure S1A), the ability to activate LRH-1 is associated with coordinated motions arising from these interactions.

### Coupled motion induced by active ligands extends to AFB

The shortest distance data shows a relationship between ligand activity and coupled motion between the ligand and helix 7. Using community analysis, we find that the connectivity between active ligands and helix 7 extends beyond the binding pocket. This analysis divides protein complexes into “communities,” groups of residues that move together. Similar to the concept behind shortest distance measurements, coupled motions among residues in a community are thought to reflect a functional relationship. Representative results for LRH-1 bound to an active ligand and an inactive ligand are shown in Figure 3A, where each community is shown in a different color. Ligand atoms are also colored according to which community they belong to, and one of the most striking results from this analysis is that the ligand community differs depending on ligand classification. Consistent with the shortest distance results, active ligands belong to the helix 7 community more frequently than inactive ligands, indicating that they move together with residues in this community more than with other communities. This is the case for 100% of active ligands and only 50% of inactive ligands, p = 0.01 by Fisher’s exact test, Figure 3A-B, Figure S2). In addition, helix 7 communities are 43% larger in the presence of active ligands *versus* inactive ligands, containing 60 +/- 4 and 39 +/- 4 residues, respectively (mean +/- SEM, p = 0.005 by two-tailed, unpaired t-test, Figure 3C *left*). In almost all cases, the larger helix 7 community contains AFB (comprised of helix 6 and beta sheets 1 and 2). AFB is a part of the helix 7 community for 83% of complexes with active ligands *versus* 25% of complexes with inactive ligands (p= 0.01, Fisher’s exact test, Figure 3B).

**Figure 3.**
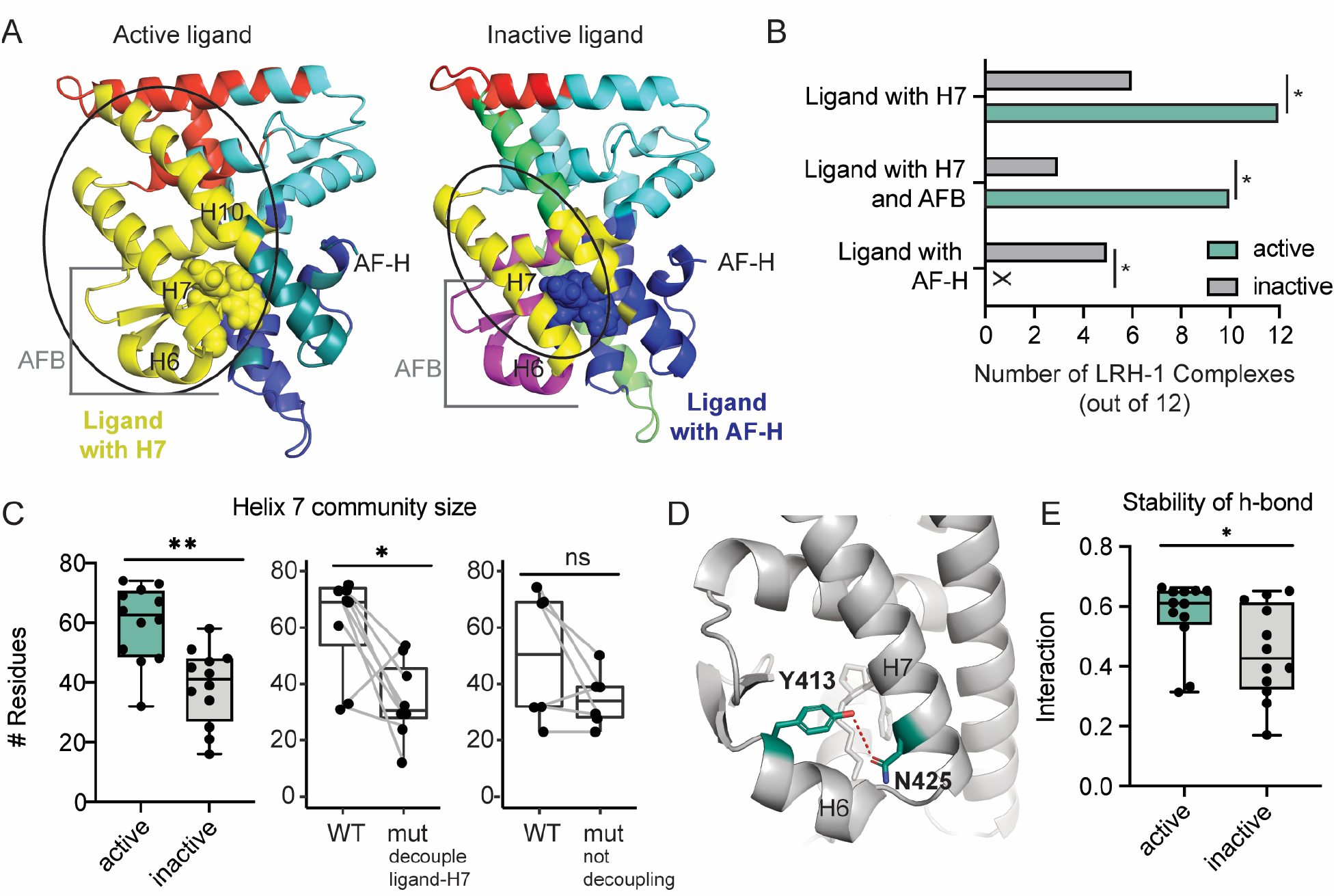
Coupled motion induced by active ligands extends to AFB. A. Representative results from community analysis for one active ligand (*left*) and one inactive ligand (*right*), mapped onto the LRH-1 ligand binding domain (PDB 4DOS). Each color represents a community, which is a group of residues that move together. Ligands are shown as spheres and colored according to the community they participate in. Complexes with active ligands tend to have larger communities containing the ligand, helix 7 and AFB (yellow, highlighted with a circle). Communities for individual complexes are shown in the supplemental material (Figure S2). B. Active ligands are more likely to belong to the community containing helix 7 than inactive ligands. Plot shows the number of complexes in which the ligand is grouped in the same community with helix 7. *, p < 0.05, Fisher’s exact test. C, *left*. The helix 7 community is larger in the presence of active *versus* inactive ligands. Each point represents the number of residues in the helix 7 community for an LRH-1 complex (n =12). Lines in the boxes indicate the mean and whiskers the range. **, p < 0.01 by two-tailed, unpaired t-test. C, *middle and right*. Mutations that decouple helix 7-ligand motions reduce the size of the helix 7 community. Each point represents the number of residues in the helix 7 community for wild type (WT) or mutant (mut) LRH-1 bound to a specific ligand. Lines connecting points follow changes for WT versus mutant LRH-1 bound to the same ligand (*e.g*., WT LRH-1 bound to ligand 5N is connected to M345L LRH-1 bound to 5N). *, p < 0.05 by two-tailed, paired t-test. D. A hydrogen bond between Y413 and N425 (shown in teal sticks) between helix 6 and helix 7. E. The hydrogen bond in *D* is more stable when LRH-1 is bound to active ligands. Each point represents an LRH-1-ligand complex, plotted as in *C*. *, p < 0.05 by two-tailed, unpaired t-test. H, helix.

We observed a hydrogen bond between residues Y413 and N425, bridging helices 6 and 7 (Figure 3D), which is more stable when active *versus* inactive ligands are bound (Figure 3E) and investigated whether this bond is important for linking motions of AFB and helix 7. Introducing an N425A mutation to a subset of active ligand complexes does not significantly decrease the size of the helix 7 community, change the participation of the ligand in it, nor does it split AFB and 7 into different communities (Figure S3). Moreover, N425A does not significantly affect ligand coupling with helix 7 in shortest distance analysis (p = 0.29 by two-tailed, paired t-test). This suggests that the increased stability of the hydrogen bond is a product of the connectivity in this region rather than the cause of it. However, mutations that decouple ligand-helix 7 motions (identified in Figure 2C) reduce the size of the helix 7 community, making it similar in size to complexes with inactive ligands (Figure 3C, *middle*). Mutations to ligand-contacting residues that do not decouple ligand-helix 7 motions do not change the community size to a level of statistical significance (Figure 3C, *right*). This supports the idea that ligand coupling with helix 7 induces connectivity between helix 7 and a larger region of the receptor.

### Inactive ligands induce communication with the AF-H

The community analysis reveals a different pattern for inactive ligands compared to active ligands: inactive ligands tend to participate in the community containing the AF-H instead of helix 7 (Figure 3A-B). This is the case for five out of the 12 complexes with inactive ligands and none of the complexes with active ligands (p = 0.03, Fisher’s exact test, Figure 3B). This connection between inactive ligands and the AF-H, an important part of the AFS, implies strong communication between these distant sites. We investigated the strength of ligand-AFS communication using suboptimal paths analysis, which quantifies correlated motion of residues along paths between two distant points in a protein or complex. It considers the complex as a network, where each Cα is a node, and the connection between each node is an edge. Nodes with greater correlated motion will have stronger edges, and the sum of edges for nodes along paths traversing two points indicates the strength of communication. The strongest (“optimal”) path and a subset of the strongest suboptimal paths are thought to convey the most information between the two points. LRH-1 complexes bound to inactive ligands have more than double the number of strong paths between the ligand and the AFS than those bound to active ligands, supporting the idea that inactive ligands promote communication with the AFS (Figure 4A). The strong paths pass through helix 3 for all complexes regardless of ligand class, suggesting that this helix is an important conduit of the signal (Figure 4C, blue lines).

**Figure 4.**
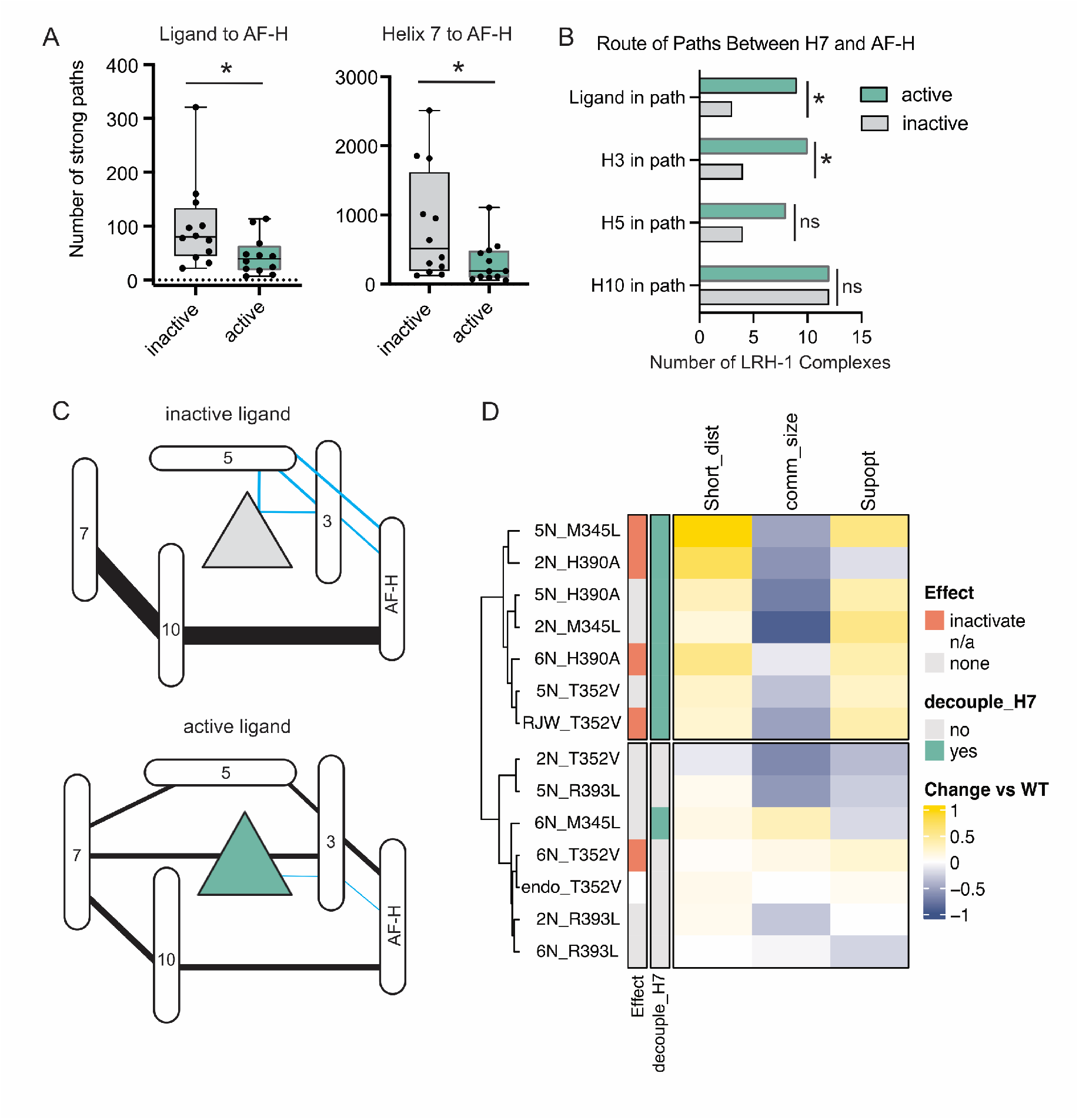
Inactive ligands induce communication with the AF-H. A. Quantification of the number of strong suboptimal paths between each ligand and the AF-H (*left*), or between helix 7 and the AF-H (*right*). Each point is a protein-ligand complex, the middle line in the boxes indicates the mean, and the whiskers indicate the range. *, p < 0.05 by a two-tailed, unpaired t-test. B. Quantification of the numbers of complexes (out of 12) in which the strong paths traverse different parts of the receptor (as indicated in the figure panel). *, p < 0.05 by Fisher’s exact test. C. Summary of the routes and strengths of communication to the AF-H originating from helix 7 (black lines) or the ligand (blue lines) in complexes containing inactive (*left*) or active (*right*) ligands. Oblong shapes represent LRH-1 helices (labeled with number or name), and triangles represent ligands. Lines were drawn between components if greater than 50% of complexes had strong suboptimal paths between them. Thickness of the lines are proportional to the mean number of strong suboptimal paths, where thicker lines indicate stronger communication. D. Heatmap summarizing the effect of LRH-1 mutations on three features identified in our analysis as being different between active and inactive ligands (1) coupled motion between helix 7 and the ligand (short_dist), (2) size of the helix 7 community (comm_size), and (3) strength of communication between helix 7 and the AFS (subopt). The color of the cells shows how these values change for mutant versus WT LRH-1 in each LRH-1-ligand complex (normalized to the range of each column to put values on the same scale). The left annotation includes the ligand/mutation combination shown in each row. The top cluster shows enrichment for a set of mutations that decouple ligand-helix7, decrease the helix 7 community size, and increase helix7-AFS communication.

To identify additional mechanisms of intramolecular communication associated with ligand activity, we examined suboptimal path strengths between several other parts of the receptor to the AFS. Communication between helix 7 and the AFS was the only comparison tested that was statistically significantly different for inactive *versus* active ligands (Table S1). Inactive ligands promote stronger helix 7-AFS communication than active ligands, with an average of nearly three times the number of strong paths (Figure 4A). Helix7-AFS communication follows different routes depending on ligand class. For inactive ligands, it mainly traverses helix 10, while active ligands induce connectivity with other parts of the receptor in addition to helix 10 (*i.e*., helix 3, helix 5, and the ligand, Figure 4B-C). Mutations that decouple ligand-helix 7 motion, including inactivating mutations, increase communication between helix 7 and AFS compared to WT LRH-1 bound to the same ligands (Figure 4D).

Together, our analysis indicates that inactive ligands are more strongly connected with the AFS via signaling pathways that originate from both the ligand binding pocket and helix 7. In contrast, active ligands have relatively weaker communication with the AFS and are instead more connected with helix 7/AFB (Figure 2-3). These results suggest that ligands communicate their identity to the AFS through intramolecular signaling networks centered on helix 7.

### The Tif2 coregulator strengthens allosteric signaling and destabilizes the AFS when LRH-1 is unliganded

NR activation involves an interplay between ligand- and coregulator-derived signals. To investigate coregulator-derived signaling, we conducted MD simulations with a fragment of Tif2 bound at the AFS. Tif2 is a coactivator and would be expected to be preferentially recruited to complexes containing active ligands. However, it also binds LRH-1 when bound to weak agonists and when unliganded (20), making it well-suited for these studies.

We first examined how Tif2 affects unliganded LRH-1 (apo-LRH-1). In absence of Tif2, apo-LRH-1 has even stronger helix 7-AFS communication than complexes containing inactive ligands (1113 strong paths *versus* 846 +/- 232 for inactive ligands, mean +/- SEM, Table S2). Tif2 binding further strengthens this communication, as well as communication to AFB, by around three-fold (Figure 5A). Additionally, Tif2 destabilizes components of the AFS (AF-H and helix 3), as measured by root mean square fluctuation (RMSF, Figure 5B-C). Therefore, Tif2 binding the apo receptor strengthens an inactive-like allostery while destabilizing the Tif2 binding site. This response may represent a mechanism for recognizing aberrant coregulator recruitment in absence of ligand.

**Figure 5.**
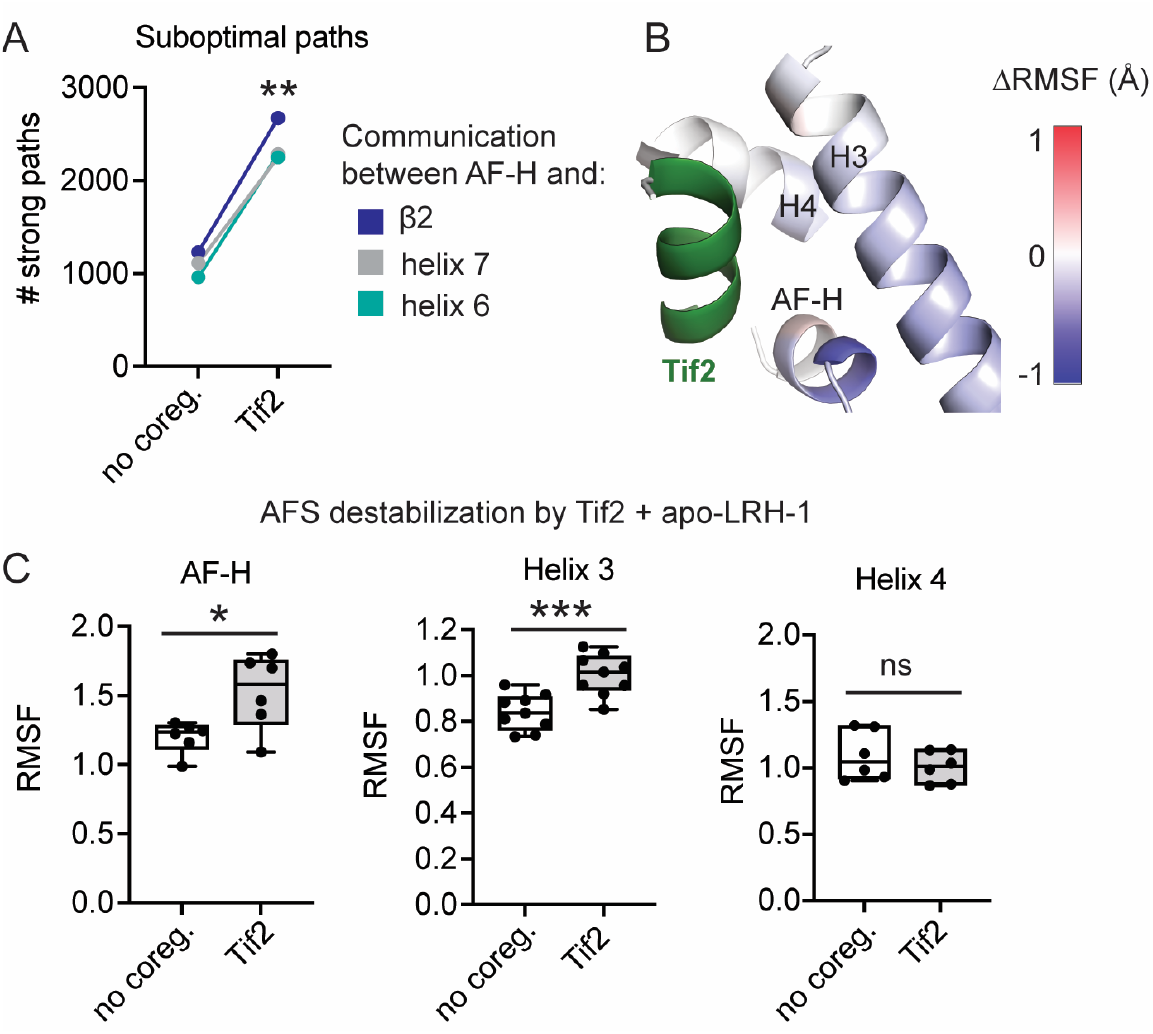
Tif2 coregulator binding apo-LRH-1 strengthens allosteric signaling and destabilizes the AFS. A. Number of strong suboptimal paths between the AF-H and helix 6 or β2 (parts of AFB) and between the AF-H and helix 7 in the presence and absence of Tif2. **, p < 0.01 by a two-tailed, paired t-test. B-C. Tif2 destabilizes the AFS when LRH-1 is unliganded. B. RMSF differences (ΔRMSF) for apo-LRH-1 with no coregulator bound minus apo-LRH-1 + Tif2 mapped onto the AFS (comprised of residues in helix 3, helix 4, and the AF-H). Negative values indicate destabilization upon Tif2 binding. C. RMSF of residues located in the AFS, either in the AF-H (*left*), helix 3 (*middle*) or helix 4 (*right*), in the presence or absence of Tif2. Each point is the RMSF of a residue in the AFS, lines in the boxes represent the mean, and whiskers represent the range. *, p < 0.05; ***, p < 0.001 by two-tailed, unpaired t-tests.

### Co-binding of a coactivator and an inactive ligand is destabilizing

For ligand-bound LRH-1, the effect of Tif2 binding differs from apo-LRH-1. For complexes with inactive ligands, Tif2 attenuates helix 7-AFS communication to a level similar to active ligand complexes (Table S2). This is the opposite effect of Tif2 binding apo-LRH-1 (Figure 5A). In complexes with active ligands, Tif2 binding does not affect the connectivity between ligands and helix 7 (by shortest distance analysis) or communication with the AFS (by suboptimal paths analysis). We did not detect differences in communication strengths between AFS and the ligand, helix 6, or β2 upon Tif2 binding for either ligand class, nor was strength of communication originating from Tif2 significantly different for active and inactive ligands (Table S1).

To better understand the effects of Tif2 on LRH-1 dynamics in the presence of different ligands, we used principal component analysis (PCA) to identify the essential motions of each complex. We obtained the top four principal components (PCs) for each complex and calculated how much they overlap by computing the root mean square inner product for each pair of complexes. The degree of overlap of the PCs indicates how similar the essential motions are for the pair. Both in the presence and absence of Tif2, the PC overlap matrices do not cluster by ligand class, size of the R^1^ group, stereochemistry, or any previously measured biological parameter (Figure 6A-B). However, PCA fluctuation analysis, which calculates the contribution of each residue to the top four PCs, revealed an important role for AFB in the essential motions of complexes containing inactive ligands when Tif2 is bound (Figure 6C-E). Therefore, while the overall PCs of the complexes do not segregate by ligand type, the per-residue decompositions in the PC fluctuation analysis reveals a common involvement of AFB in complexes with inactive ligands.

**Figure 6.**
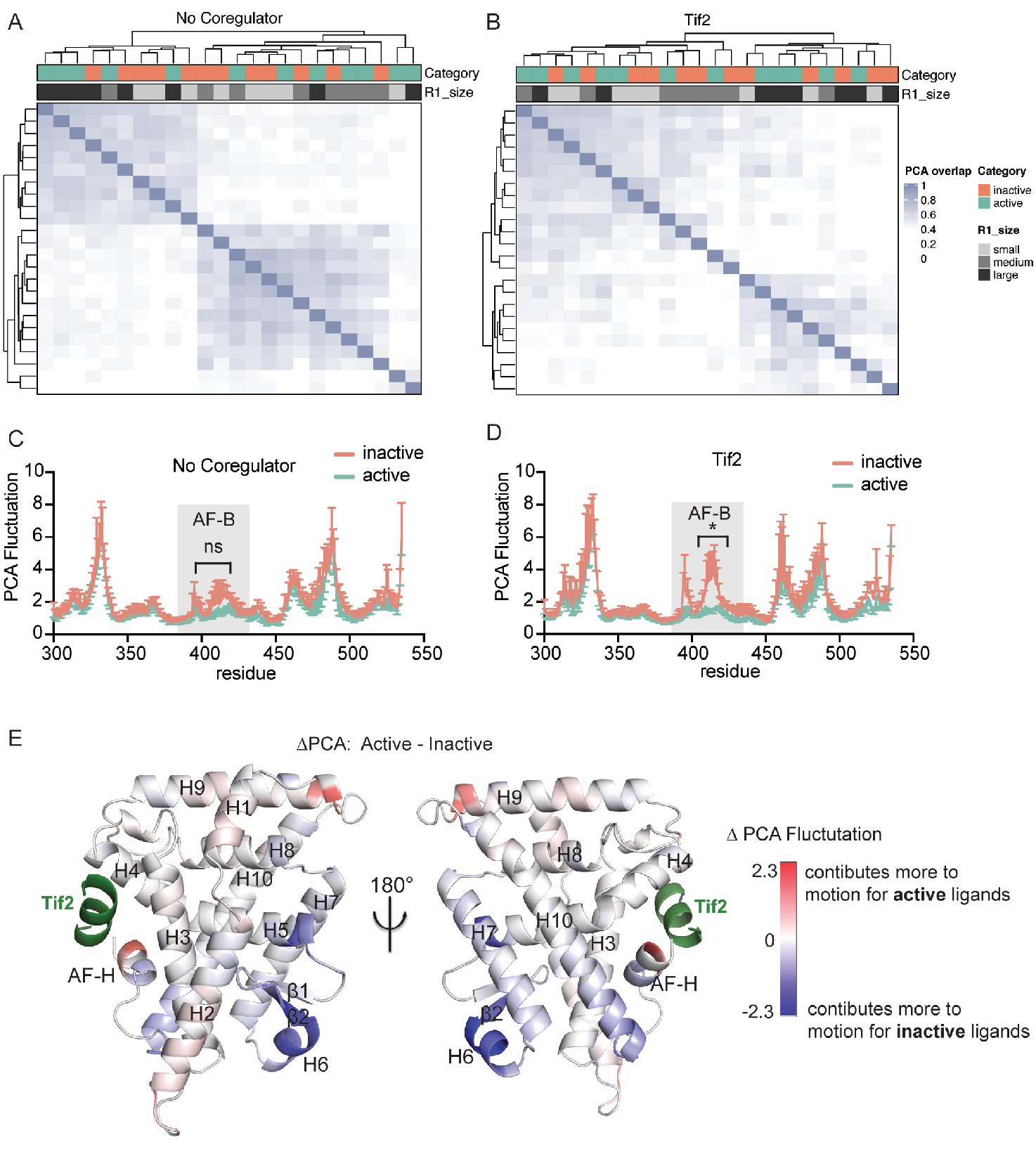
PCA shows the importance of AFB for inactive ligands. A-B. The top four PCs for each proteinligand complex were determined, and the extent of overlap between these PCs is shown for each complex in the absence (*A*) or presence (*B*) of Tif2. C-D. PCA fluctuation plots. Each line represents the median value; error bars are the 95% CI. *, p < 0.05 by two-way ANOVA followed by Sidak’s multiple comparisons test. Shaded areas AF-B. E. Difference in PCA fluctuations for active minus inactive ligands in the presence of Tif2 mapped onto the LRH-1 structure.

RMSF analysis supports the PCA results, indicating that AFB motion is a major feature discriminating active and inactive ligands when Tif2 is bound. In absence of Tif2, all of the ligands are stabilizing relative to apo-LRH-1, with the strongest effect on flexible regions near the ligand binding pocket (*e.g*. parts of helix 2 and AFB, Figure 7A-B). There are no statistically significant differences between active and inactive ligands in absence of Tif2 (Figure 7A), but Tif2 binding differentially affects portions of AFB, the AF-H, and the pre-AFH loop (Figure 7C-D). The differences arise from increased fluctuations with inactive ligands rather than from stabilization by active ligands (Figure 7E). Fluctuations induced by inactive ligands + Tif2 are similar in magnitude to that of apo-LRH-1 both with and without Tif2 bound, Figure 7A; 7C). This suggests that, while all ligands tend to stabilize AFB, inactive ligands are unable to maintain the stabilization when Tif2 is bound. These findings provide insight into the PCA results, suggesting that the essential motions of inactive ligand complexes are driven by fluctuations at AFB rather than coordinated motions at this site.

**Figure 7.**
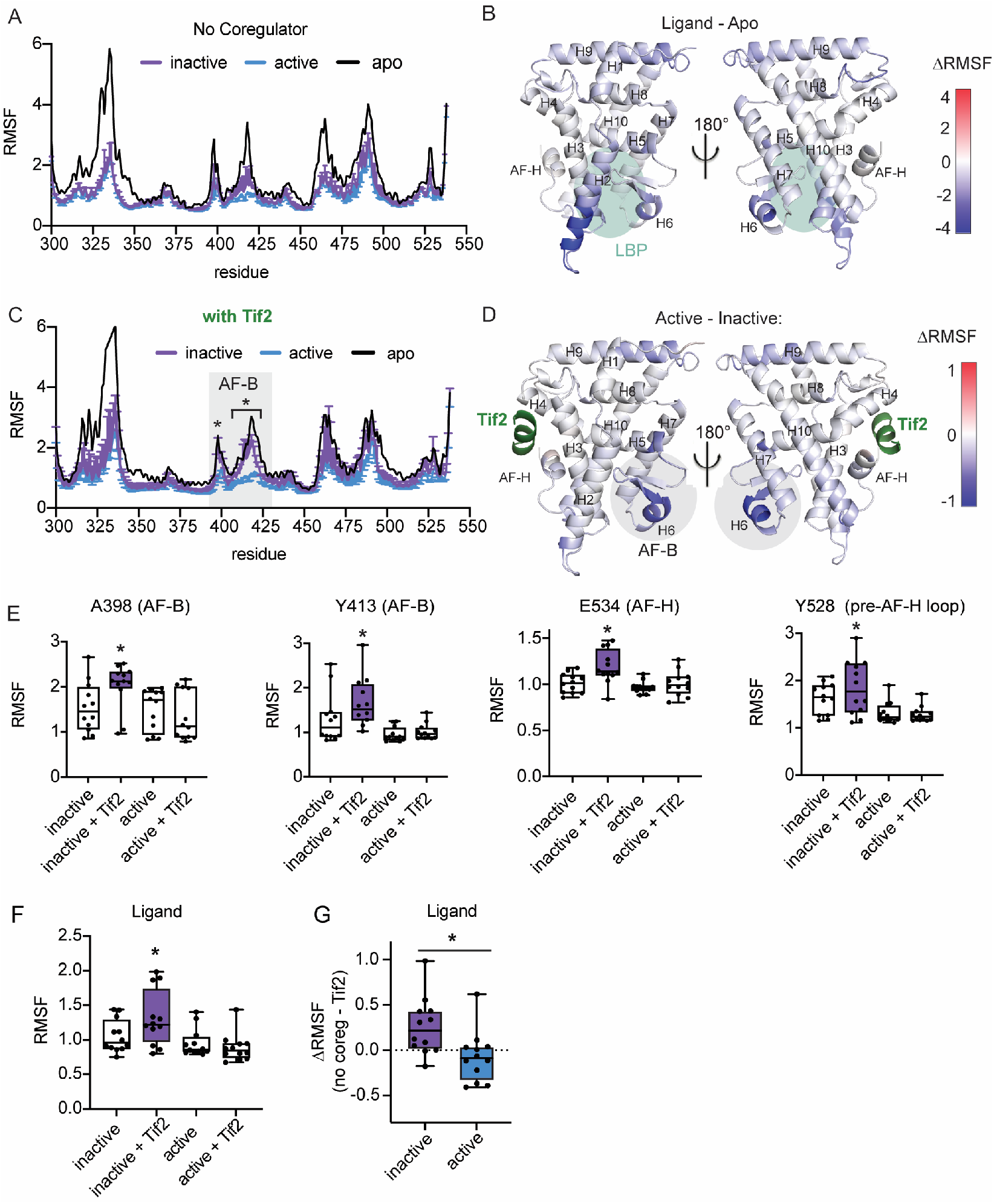
Co-binding of a coactivator and an inactive ligand is destabilizing. A. RMSF for each LRH-1 residue when apo or bound to active or inactive ligands when no coregulator bound. Median RMSF values +/- 95% CI are shown for ligand-bound complexes (n = 12). B. The difference in RMSF (ΔRMSF) of ligand-bound LRH-1 (n = 24) minus RMSF of apo-LRH-1 is mapped onto the LRH-1 structure to show the regions affected the most by ligand binding. Teal shaded area indicates the ligand binding pocket (LBP). The color of helix 2 (H2) in the right panel appears lighter than in the left because it is in the background after rotation of the complex. C. RMSF plot comparing active *versus* inactive ligands (n = 12) in the presence of Tif2. Each line is the median RMSF, and the error bars represent the 95% CI. *, p < 0.05 by two-way ANOVA followed by Sidak’s multiple comparisons test. The RMSF plot of apo-LRH-1 bound to Tif2 is shown for comparison. D. ΔRMSF of active minus inactive ligands in the presence of Tif2 (shown in green), mapped onto the LRH-1 structure. Grey shaded areas in *C* and *D* indicate AFB. E. Plots of RMSF values of the indicated residues, showing destabilization with the combination of inactive ligands and Tif2. Midpoints represent the median, error bars are the range. *, p < 0.05 *versus* the other groups by one-way ANOVA followed by Dunnett’s multiple comparisons test. F. Ligand RMSF values showing that the combination of inactive ligands with Tif2 destabilizes the ligand. Plots and statistics are done the same way as panel *E*. G. The effect of Tif2 on ligand fluctuations for individual LRH-1-ligand complexes was determined by calculating ΔRMSF (RMSF with no coregulator minus RMSF+Tif2). A value of zero would mean no change upon Tif2 binding, negative values indicate stabilization, and positive values indicate destabilization. *, p < 0.05 by two-tailed, paired t-tests.

Ligand fluctuations (RMSF calculated over ligand atoms) mirror the distinctive pattern of AFB fluctuations on a smaller scale. Tif2 destabilizes inactive ligands (Figure 7F) and has the opposite effect on active ligands, which can be seen by comparing how RMSF changes for each complex (ΔRMSF, Figure 7G). The increased ligand and receptor fluctuations could explain how communication between the AFS and helix 7 is attenuated with the combination of an inactive ligand and Tif2 (Table S2) and may reflect a disharmony arising from the combination of an inactive ligand with a coactivator.

## Discussion

NRs are attractive drug targets, capable of transcribing varied gene programs depending on ligand-induced coregulator associations. These studies provide insights into NR regulation by identifying how LRH-1 ligands transmit their identities to the site of coregulator binding at the AFS. Analyses of a large set of closely related compounds with established activity profiles enabled the discovery of distinct intramolecular signaling circuits utilized by active and inactive ligands (Figure 8). When LRH-1 is unliganded or bound to inactive ligands, there is strong communication with the AFS, mediated by the ligand and helix 7 (shown with suboptimal paths analysis, Figures 4-5). Activating ligands reduce helix 7-AFS communication and induce connectivity with AFB (shown with shortest distance and community analyses, Figures 2-3). Helix 7 is at the center of both circuits, sensing the ligands and changing its motion to direct the signaling toward either AFB or the AFS. These results could inform the design of LRH-1 modulators, which are sought as treatments for metabolic diseases, cancers, and inflammatory bowel diseases. For example, modifications that make polar interactions with helix 7 residues could increase receptor activation by increasing ligand-helix 7 coupling.

**Figure 8.**
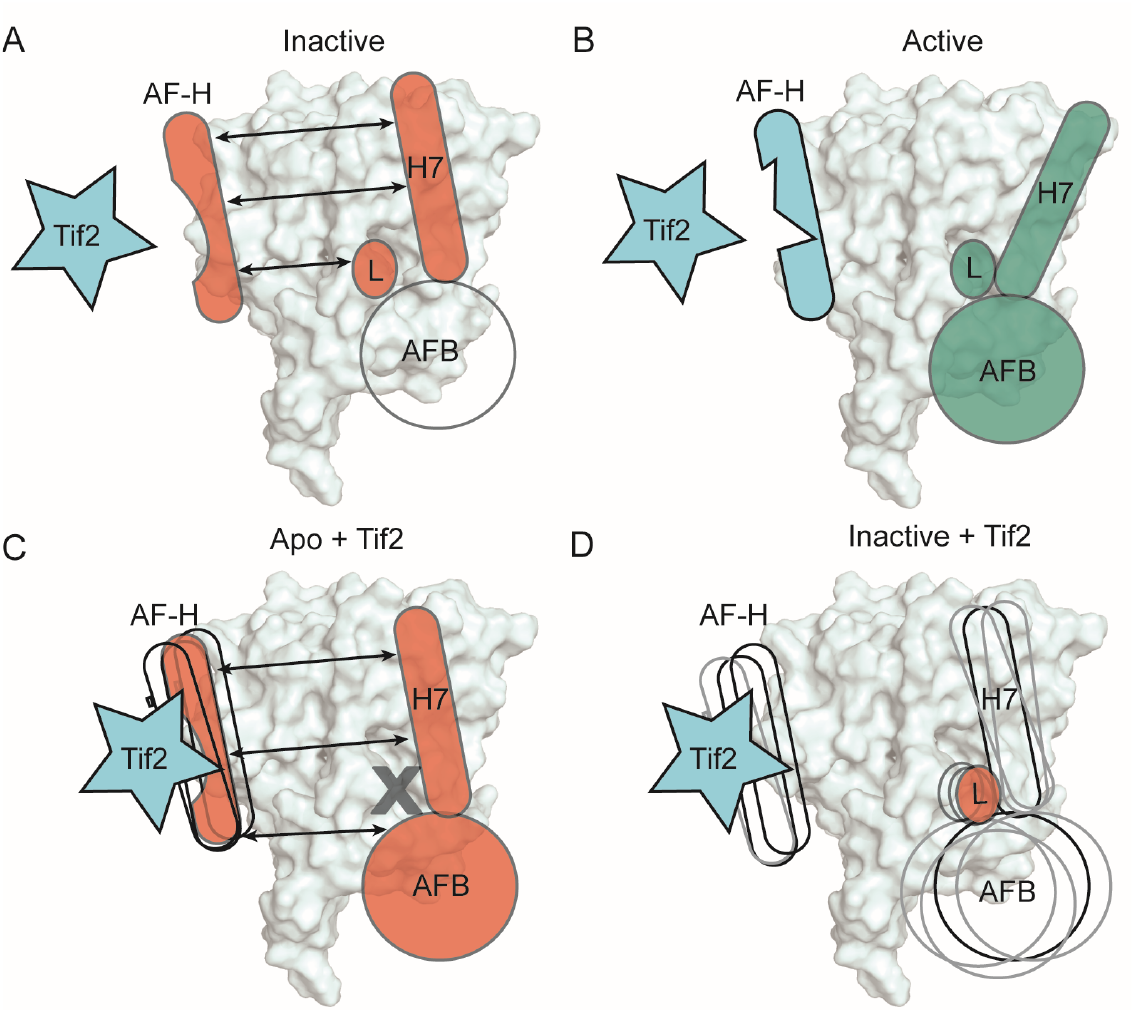
Proposed model: LRH-1 tunes allosteric signaling in response to ligands and ligand-coregulator combinations. A. Inactive ligands communicate with the AF-H and promote communication between helix 7 and the AF-H. B. Active ligands communicate with helix 7 and AFB and do not signal strongly to the AF-H. C. Tif2 binding apo-LRH-1 destabilizes the AF-H and strengthens allosteric communication. D. Tif2 binding together with an inactive ligand destabilizes the ligand, AFB, and the AFS.

We discovered the importance of helix 7 using an unbiased clustering of shortest distance data to identify ligand contacts associated with LRH-1 activation in cellular assays (Figure 2). Since the synthetic design of the 6HP ligands used in this study aimed to strengthen interactions with helix 3, it is not surprising that helix 3 interactions were identified (Figure 2A, C). The involvement of helix 7 was unexpected, since the ligands interact with it using their identical hydrophobic “tails” (Figure 1B). Studies probing LRH-1 dynamics in solution have hinted that the motion of the N-terminus of helix 7 is sensitive to ligand status (22,24,26), even with a phospholipid agonist that does not share the 6HP scaffold (20). However, we might have overlooked its importance if not for the unbiased approach to the MD analysis. Our data show that interactions with helix 3 can affect ligand coupling with helix 7, which may explain how these ligands have differential effects on helix 7. Inactivating mutations in helix 3 disrupt the coupled motion of the ligands with helix 7 (Figure 2C). Additionally, routes of communication between helix 7 and the AF-H preferentially pass through helix 3 in the presence of active ligands (Figure 4B).

A striking aspect of the signaling mechanism is the strong AFS communication by inactive ligands or in apo-LRH-1. The purpose of this signaling is unclear, but we hypothesize that this is used to avoid premature turnover or aberrant activation of LRH-1 in the nucleus. In contrast to type I NRs (*e.g*. steroid receptors), which are localized in the cytosol and stabilized by chaperone proteins prior to ligand binding, LRH-1 belongs to a subset of NRs that reside in the nucleus (19). Persistent communication of the apo state may have evolved as a way for NRs to indicate they are well-folded and available for ligand binding, avoiding degradation without needing to be stabilized by chaperones. Additionally, NRs residing in the nucleus often have lipophilic binding pockets and must recognize specific lipid ligands among similar, more abundant endogenous lipids. For example, PPAR family members are activated by fatty acids with specific chain lengths, and LRH-1 binds many types of phospholipids that do not activate it (11). This binding promiscuity suggests the need for a barrier or checkpoint to avoid improper activation by nonspecific ligands. In the case of LRH-1, strong AFS communication could serve as the barrier, which is overcome by ligands that induce sufficient coupling with helix 7. This possibility will be interesting to study further when more is known about endogenous LRH-1 ligands. The natural LRH-1 ligands are likely phospholipids, but we currently lack the depth of structure-function data to conduct a similar study with phospholipid ligands.

This work also provides insights into allosteric signaling by the Tif2 coactivator. When LRH-1 is unliganded, Tif2 increases the strength of helix 7-AFS signaling (Figure 5). This suggests that Tif2 recognizes the apo receptor using a helix 7-AFS signaling network like inactive ligands. However, there are important distinctions in how Tif2 affects LRH-1 dynamics when it is apo or bound to inactive ligands. Tif2 increases helix7-AFS signaling in apo-LRH-1, but it has the opposite effect in the presence of inactive ligands. In addition, Tif2 destabilizes LRH-1 when bound to inactive ligands, increasing ligand and receptor fluctuations compared to complexes containing active ligands. The destabilizing effect is most prominent in AFB, where fluctuations dominate the complexes and define their principal motions (Figures 6-7). These fluctuations likely reflect a “frustrated” state in which a coactivator is paired with an inappropriate ligand. Therefore, we have identified dynamical changes associated with multiple bound states of the receptor, mediated by both coordinated residue motions and by fluctuations (Figure 8). Activating ligands induce coordinated, productive motions with residues in helix 7 and AFB, attenuating signaling to the AF-H. Tif2 communicates via coordinated motions or destabilizing fluctuations depending on ligand status. Both types of allosteric communication have been reported in other systems (30–35).

Together, these studies have revealed novel features of LRH-1 allostery, but they do not describe all possible receptor conformations and allosteric circuits. LRH-1 has not been crystallized bound to an antagonist or an inactive ligand, and its true “inactive” conformation is unknown. Therefore, the MD simulations described here are modeling the response of placing an inactive ligand in the context of an active receptor rather than a transition to an inactive state. The conformational dynamics of apo-LRH-1 are also incompletely understood. Apo-LRH-1 has been crystallized with two coregulators, and both structures depict the AFS in the canonical “active” conformation. However, biophysical assays demonstrate it has conformational flexibility to interact with coregulators that require large displacements of the AF-H. Finally, NRs interact with many coregulators, which likely utilize different allosteric mechanisms. Tif2 has been crystallized with many LRH-1-ligand complexes, and it interacts with LRH-1 in the presence of both strong and weak agonists and when the receptor is unliganded (20,21,24–26,29,36,37). This makes Tif2 well-suited for studying allosteric signaling by ligands with a wide range of activities. Investigating conformational changes and allostery associated with other coregulator-ligand combinations will be important in future studies.

## Materials and methods

### Structure Preparation

A total of 24 ligands were used in MD simulations. When possible, LRH-1-ligand complexes were prepared directly from crystal structures (PBD IDs 5L11, 5SZY, 6OR1, 6OQX, 6OQY containing ligands RJW100, Endo, 2N, 5N, and 6N, respectively). All other ligand complexes were constructed by manually modifying atoms of ligands in the structure containing the most similar ligand (Table 1). The model of LRH-1 in the apo state bound to the Tif2 coregulator peptide was prepared from PBD 4PLD. All models contained LRH-1 residues 300-583 (the ligand binding domain) and residues 742-751 of Tif2 (the NR box 3 domain that interacts with LRH-1). For each complex, non-Tif2-bound complexes were also constructed for simulation by removing Tif2 peptide.

Five LRH-1 mutants were explored for a subset of the complexes: M345L, H390A, T352V, R393L and N425A (Table 1). Mutants complexes were created by introducing mutations *in silico* using x-leap in AmberTools18(38) or in Coot(39). In total, 69 models were used for MD simulations (Table S1). *MD simulations*. Molecular dynamics simulations were performed on the 69 complexes described above. All LRH-1 complexes were solvated with TIP3P water(40) in a truncated octahedron, with a 10 Å water buffer surrounding the protein complex. Na+ and Cl-ions were added to neutralize the protein and achieve physiological salt conditions. The xleap tool of AmberTools18(38) was used to generate all systems for MD simulations. The parm99-bsc0 forcefield(41) was used for proteins while the Generalized Amber Forcefield (GAFF)(42,43) was used in Antechamber(44) to parameterize all synthetic agonists. Minimizations and MD simulations were performed in Amber18. All minimizations were performed using 5000 steps of steepest descent followed by 5000 steps of conjugate gradient. The following series of positional restraints were used in sequential minimizations: i) 500 kcal/mol·Å^2^ on all protein and ligand atoms; ii) 100 kcal/mol·Å^2^ on all protein and ligand atoms, iii) 100 kcal/mol·Å^2^ on ligand atoms only, iv) no restraints. Minimized systems were heated from 0 to 300 K with a 100-ps run, constant volume periodic boundaries and 5 kcal/mol·Å^2^ restraints on all protein and ligand atoms. Equilibration and production runs were performed using AMBER on GPUs(45,46). Runs were performed in triplicate with the following protocol: i) 10-ns run with 10 kcal/mol·Å^2^ restraints on all solute atoms, ii) 10-ns run with 1 kcal/mol·Å^2^ restraints on solute atoms, iii) 10-ns run with 1 kcal/mol·Å^2^ restraints on ligand atoms alone, and finally iv) 500 ns production trajectories with no restraints on any atoms in the NPT ensemble. All bonds between heavy atoms and hydrogens were fixed with the SHAKE algorithm (47). A cutoff distance of 10 Å was used to evaluate long-range electrostatics with particle mesh Ewald and for van der Waals forces.

### Analysis of MD simulations

Structural averaging was performed using the CPPTRAJ(48) module of AmberTools18. The ‘strip’ and ‘trajout’ commands of CPPTRAJ were used to remove solvent atoms and obtain twenty-five thousand evenly spaced frames from each simulation for analysis. For each complex, triplicate runs were combined to yield seventy-five thousand frames. Root mean square fluctuation analysis was performed on Cα atoms of protein residues and computed for each frame in the trajectory relative to the initial structure.

Cartesian covariances were calculated for each complex in Carma(49). The NetworkView plugin in VMD(50) was used to produce dynamic networks for each system. Briefly, networks were constructed by defining each protein residue as a node and using distance between nodes over the simulation to define contacts/ ‘edges.’ If heavy atoms of one node were within 4.5 Å of heavy atoms from another non-neighboring node for 75% of the trajectory, an edge is created between the nodes. Edges were not created between neighboring nodes. Calculated covariances were used to weight edges of dynamic networks, such that weights are inversely proportional to the calculated pairwise correlation between the nodes. The protocol described in(51) was followed.

To describe communication between different LRH-1 sites, suboptimal paths were generated from dynamic networks using the Floyd-Warshall algorithm(53). Briefly, suboptimal paths are chains of edges that predict route and strength of communication between two points. For all paths (i.e. chains of edges) connecting two points, the optimal path is the one for which the sum of edges is the lowest. Correspondingly, correlation is highest between these nodes. Therefore, a collection of shortest suboptimal paths conveys the strongest communication between two distant nodes (i.e. source and sink nodes), also revealing the direction of signaling. We obtain the set of ‘strong’ suboptimal paths by adding a cutoff (75) to the path length of the optimal path and extracting all paths that lie within this range. For source and sink nodes, we used LRH-1 residues 404 (β2), 419 (helix 6), 425 (helix 7), 534 (AF-H), and Tif2 residue 748.

To compare the strength of communication in a pair of nodes across multiple complexes, we defined the parameter ‘shortest distance.’ If the pair of nodes is linked by an edge, shortest distance is the weight of the edge as determined in the dynamic network. If the pair of nodes is not linked by an edge, shortest distance is the length of the optimal path linking the nodes.

To reveal the most important motions in LRH-1 ligand complexes, a principal component analysis (PCA) was performed over the backbone atoms using CPPTRAJ. Correlation matrices were determined for the protein monomers over the simulations and then diagonalized to obtain eigenvectors and eigenvalues of the system. Each eigenvector describes a dimension of positional fluctuations in the system and is weighted by its corresponding eigenvalue. A projection of the trajectory onto each eigenvector is a principal component. Typically, displacements along the top 3-5 eigenvectors describe the majority of the motions observed over a simulation. To determine the contributions by individual residues to these important motions, we performed a per-residue decomposition of the positional fluctuations represented in the top 4 principal components. For each complex, we report the sum of these decompositions as PCA fluctuations as follows:

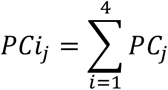

Where j = protein residue, i = principal component

To identify groups of nodes with correlated motions, communities were generated from dynamic networks using the Girvan-Newman algorithm(52). The minimum number of communities possible was generated while maintaining at least 98% modularity. The “helix 7 community” was defined as containing the residues in helix 7 contacting the ligands (L424, L427, M428, A431). AFB was defined as residues 401-421. The “AF-H community” was defined as the community containing residue L532.

### Statistics and data visualization

Heatmaps, matrices, and graphs were made using the R packages ComplexHeatmap, tidyverse, ggplot2, and ggpubr (R version 3.6.1)(54–58). Some graphs were made using GraphPad Prism, v.9. Mapping of RMSF and PCA onto structures was done in Pymol (v.2.1.1), replacing b factors with the median ΔRMSF or ΔPCA and then coloring the receptors by the replaced values. Statistics were conducted with GraphPad Prism or R, and the statistical tests used are indicated in the figure legends.

## Data Availability

Data in the manuscript with be shared upon request by the corresponding authors, Eric Ortlund or Denise Okafor

## Supporting Information

This article contains supporting information.

## Acknowledgements

none

## Funding and additional information

This work was supported by a Burroughs Wellcome Fund award to C.D.O and award number R01DK114213 and R01DK114213 -01S1 from the National Institutes of Health and an Emory Catalyst Award to E.A.O. The content is solely the responsibility of the authors and does not necessarily represent the official views of the National Institutes of Health.

## Abbreviations

6HP: hexahydropentelene
AFS: activation function surface
AFB: activation function B
AF-H: activation function helix
DSF: differential scanning fluorimetry
LRH-1: liver receptor homolog-1
MD: molecular dynamics
NR: nuclear receptor
PC: principal component
PCA: principal component analysis
RMSF: root mean square fluctuation
Tif2: transcriptional intermediary factor 2
T_m_: melting temperature

## References

1. Tobin, J. F., and Freedman, L. P. (2006) Nuclear receptors as drug targets in metabolic diseases: new approaches to therapy. Trends Endocrinol Metab 17, 284–290

2. Kojetin, D. J., and Burris, T. P. (2014) REV-ERB and ROR nuclear receptors as drug targets. Nature Reviews Drug Discovery 13, 197–216

3. Sladek, F. (2003) Nuclear receptors as drug targets: new developments in coregulators, orphan receptors and major therapeutic areas. Expert Opin Ther Targets 7, 679–684

4. Zhao, L., Zhou, S., and Gustafsson, J.-A. (2019) Nuclear Receptors: Recent Drug Discovery for Cancer Therapies. Endocrine Reviews 1, 1207–1249

5. Cave, M. C., Clair, H. B., Hardesty, J. E., Falkner, K. C., Feng, W., Clark, B. J., Sidey, J., Shi, H., Aqel, B. A., McClain, C. J., and Prough, R. A. (2016) Nuclear receptors and nonalcoholic fatty liver disease Biochim Biophys Acta 1859, 1083–1099

6. Löwenberg, M., Stahn, C., Hommes, D. W., and Buttgerit, F. (2006) Novel insights into mechanisms of glucocorticoid action and the development of new glucocorticoid receptor ligands. Steroids 73, 1025–1029

7. Hardy, R. S., Raza, K., and Cooper, M. S. (2020) Therapeutic glucocorticoids: mechanisms of actions in rheumatic diseases. Nature Reviews Rheumatology 16, 133–144

8. Jordan, V. C., and Brodie, A. M. H. (2007) Development and evolution of therapies targeted to the estrogen receptor for the treatment and prevention of breast cancer. Steroids 72, 7–25

9. Coste, A., Dubuquoy, L., Barnouin, R., Annicotte, J.-S., Magnier, B., Notti, M., Corazza, N., Antal, M. C., Metzger, D., Desreumaux, P., Brunner, T., Auwerx, J., and Schoonjans, K. (2007) LRH-1-mediated glucocorticoid synthesis in enterocytes protects against inflammatory bowel disease. Proc Natl Acad Sci U S A 104, 13098–13103

10. Bayrer, J. R., Wang, H., Nattiv, R., Suzawa, M., Escusa, H. S., Fletterick, R. J., Klein, O. D., Moore, D. D., and Ingraham, H. A. (2018) LRH-1 mitigates intestinal inflammatory disease by maintaining epithelial homeostasis and cell survival. Nat Commun 9, 4055

11. Lee, J. M., Lee, Y. K., Mamrosh, J. L., Busby, S. A., Griffin, P. R., Pathak, M. C., Ortlund, E. A., and Moore, D. D. (2011) A nuclear-receptor-dependent phosphatidylcholine pathway with antidiabetic effects. Nature 474, 506–510

12. Nadolny, C., and Dong, X. (2015) Liver receptor homolog-1 (LRH-1): a potential therapeutic target for cancer. Cancer Biol Ther 16, 997–1004

13. Lu, T., Makishima, M., JJ, R., Schoonjans, K., Kerr, T., Auwerx, J., and DJ, M. (2000) Molecular Basis for Feedback Regulation of Bile Acid Synthesis by Nuclear Receptors. Mol Cell 6, 507–515

14. Venteclef, N., Smith, J. C., Goodwin, B., and Delerive, P. (2006) Liver receptor homolog 1 is a negative regulator of the hepatic acute-phase response. Mol Cell Biol 26, 6799–6807

15. Out, C., Hageman, J., Bloks, V. W., Gerrits, H., Sollewijn Gelpke, M. D., Bos, T., Havinga, R., Smit, M. J., Kuipers, F., and Groen, A. K. (2011) Liver receptor homolog-1 is critical for adequate up-regulation of Cyp7a1 gene transcription and bile salt synthesis during bile salt sequestration. Hepatology 53, 2075–2085

16. Wagner, M., Choi, S., Panzitt, K., Mamrosh, J. L., Lee, J. M., Zaufel, A., Xiao, R., Wooton-Kee, R., Stahlman, M., Newgard, C. B., Boren, J., and Moore, D. D. (2016) Liver receptor homolog-1 is a critical determinant of methyl-pool metabolism. Hepatology 63, 95–106

17. Bolado-Carrancio, A., Riancho, J. A., Sainz, J., and Rodriguez-Rey, J. C. (2014) Activation of nuclear receptor NR5A2 increases Glut4 expression and glucose metabolism in muscle cells. Biochem Biophys Res Commun 446, 614–619

18. Mueller, M., Cima, I., Noti, M., Fuhrer, A., Jakob, S., Dubuquoy, L., Schoonjans, K., and Brunner, T. (2006) The nuclear receptor LRH-1 critically regulates extra-adrenal glucocorticoid synthesis in the intestine. J Exp Med 203, 2057–2062

19. Weikum, E. R., Liu, X., and Ortlund, E. A. (2018) The nuclear receptor superfamily: A structural perspective. Protein Sci

20. Musille, P. M., Pathak, M. C., Lauer, J. L., Hudson, W. H., Griffin, P. R., and Ortlund, E. A. (2012) Antidiabetic phospholipid-nuclear receptor complex reveals the mechanism for phospholipid-driven gene regulation. Nat Struct Mol Biol 19, 532–537, S531–532

21. Musille, P. M., Kossmann, B. R., Kohn, J. A., Ivanov, I., and Ortlund, E. A. (2015) Unexpected Allosteric Network Contributes to LRH-1 Coregulator Selectivity. J Biol Chem 291, 1411–1426

22. Mays, S. G., Flynn, A. R., Cornelison, J. L., Okafor, C. D., Wang, H., Wang, G., Huang, X., Donaldson, H. N., Millings, E. J., Polavarapu, R., Moore, D. D., Calvert, J. W., Jui, N. T., and Ortlund, E. A. (2019) Development of the first low nanomolar liver receptor homolog-1 agonist through structure-guided design. J Med Chem 62, 11022–11034

23. D’Agostino, E. H., Flynn, A. R., Cornelison, J. L., Mays, S. G., Patel, A., Jui, N. T., and Ortlund, E. A. (2020) Development of a Versatile and Sensitive Direct Ligand Binding Assay for Human NR5A Nuclear Receptors. ACS Med Chem Lett 11, 365–370

24. Mays, S. G., Okafor, C. D., Whitby, R. J., Goswami, D., Stec, J., Flynn, A. R., Dugan, M. C., Jui, N. T., Griffin, P. R., and Ortlund, E. A. (2016) Crystal Structures of the Nuclear Receptor, Liver Receptor Homolog 1, Bound to Synthetic Agonists. J Biol Chem 291, 25281–25291

25. Cornelison, J. L., Cato, M. L., Johnson, A. M., D’Agostino, E. H., Melchers, D., Patel, A. B., Mays, S. G., Houtman, R., Ortlund, E. A., and Jui, N. T. (2020) Development of a new class of liver receptor homolog-1 (LRH-1) agonists by photoredox conjugate addition. Bioorganic & Medicinal Chemistry Letters 30, 1–18

26. Mays, S. G., Okafor, C. D., Tuntland, M. L., Whitby, R. J., Dharmarajan, V., Stec, J., Griffin, P. R., and Ortlund, E. A. (2017) Structure and Dynamics of the Liver Receptor Homolog 1-PGC1alpha Complex. Mol Pharmacol 92, 1–11

27. Mays, S. G., D’Agostino, E. H., Flynn, A. R., Huang, X., Wang, G., Liu, X., Millings, E. J., Okafor, C. D., Patel, A., Cato, M. L., Cornelison, J. L., Melchers, D., Houtman, R., Moore, D. D., Calvert, J. W., Jui, N. T., and Ortlund, E. A. (2022) A phospholipid mimetic targeting LRH-1 ameliorates colitis. Cell Chem Biol

28. Knapp, B., Ospina, L., and Deane, C. M. (2018) Avoiding False Positive Conclusions in Molecular Simulation: The Importance of Replicas. J Chem Theory Comput 14, 6127–6138

29. Whitby, R. J., Stec, J., Blind, R. D., Dixon, S., Leesnitzer, L. M., Orband-Miller, L. A., Williams, S. P., Willson, T. M., Xu, R., Zuercher, W. J., Cai, F., and Ingraham, H. A. (2011) Small molecule agonists of the orphan nuclear receptors steroidogenic factor-1 (SF-1, NR5A1) and liver receptor homologue-1 (LRH-1, NR5A2). J Med Chem 54, 2266–2281

30. Kumar, V., Hoag, H., Sader, S., Scorese, H. L., and Wu, C. (2020) GDP Release from the Open Conformation of Gα Requires Allosteric Signaling from the Agonist-Bound Human β2 Adrenergic Receptor. Journal of Chemical Information and Modeling 60, 4064–4075

31. Sogunmez, N., and Akten, E. D. (2019) Intrinsic Dynamics and Causality in Correlated Motions Unraveled in Two Distinct Inactive States of Human β2-Adrenergic Receptor. Journal of Physical Chemistry B 123, 3630–3642

32. Koehler, C., Carlstroem, G., Weininger, U., Tangefjord, S., Ullah, V., Lepisto, M., Karlsson, U., Papvoine, T., Edman, K., and Akke, M. (2020) Dynamic allosteric communication pathway directing differential activation of the glucocorticoid receptor. Science Advances 6, 1–10

33. Galdadas, I., Qu, S., Oliveira, A. S. F., Olehnovics, E., Mack, A. R., Mojica, M. F., Agarwal, P. K., Tooke, C. L., Gervasio, F. L., Spencer, J., Bonomo, R. A., Mulholland, A. J., and Haider, S. (2021) Allosteric communication in class A beta-lactamases occurs via cooperative coupling of loop dynamics. eLife 10

34. Schulze, J. O., Saladino, G., Busschots, K., Neimanis, S., Suss, E., Odadzic, D., Zeuzem, S., Hindie, V., Herbrand, A. K., Lisa, M. N., Alzari, P. M., Gervasio, F. L., and Biondi, R. M. (2016) Bidirectional Allosteric Communication between the ATP-Binding Site and the Regulatory PIF Pocket in PDK1 Protein Kinase. Cell Chem Biol 23, 1193–1205

35. Whitford, P. C., Gosavi, S., and Onuchic, J. N. (2008) Conformational transitions in adenylate kinase. Allosteric communication reduces misligation. J Biol Chem 283, 2042–2048

36. Mays, S. G., Stec, J., Liu, X., D’Agostino, E. H., Whitby, R. J., and Ortlund, E. A. (2020) Enantiomer-specific activities of an LRH-1 and SF-1 dual agonist. Sci Rep 10, 22279

37. Mays, S. G., D’Agostino, E. H., Flynn, A. R., Huang, X., Wang, G., Liu, X., Millings, E. J., Okafor, C. D., Patel, A., Cato, M. L., Cornelison, J. L., Melchers, D., Houtman, R., Moore, D. D., Calvert, J. W., Jui, N. T., and Ortlund, E. A. (2020) Tapping into a phospholipid-LRH-1 axis yields a powerful anti-inflammatory agent with in vivo activity against colitis. bioRxiv

38. Case, D., Ben-Shalom, I. Y., Brozell, S. R., Cerutti, D. S., Cheatham III, T. E., Cruzeiro, V. W. D., Darden, T. A., Duke, R. E., Ghoreishi, M. K., Gilson, H., Gohlke, H., Goetz, A. W., Greene, D., Harris, R., Homeyer, N., Izadi, S., Kovalenko, A., Kurtzman, T., Lee, T. S., LeGrand, S., Li, P., Lin, C., Liu, J., Luchko, T., Luo, R., Mermelstein, D. J., Merz, K. M., Miao, Y., Monard, G., Nguyen, C., Nguyen, H., Omelyan, I., Simmerling, C. L., Smith, J., Salomon-Ferrer, R., Swails, J., Walker, R. C., Wang, J., Wei, H., Wolf, R. M., Wu, X., Xiao, L., York, D. M., and Kollman, K. A. (2018) Amber 2018. University of California, San Franscisco

39. Emsley, P., and Cowtan, K. (2004) Coot: model-building tools for molecular graphics. Acta Crystallogr D Biol Crystallogr 60, 2126–2132

40. Jorgensen, W. L., Chandrasekhar, J., Madura, J. D., Impey, R. W., and Klein, M. L. (1983) Comparison of simple potential functions for simulating liquid water. The Journal of chemical physics 79, 926–935

41. Pérez, A., Marchán, I., Svozil, D., Sponer, J., Cheatham, T. E., Laughton, C. A., and Orozco, M. (2007) Refinement of the AMBER force field for nucleic acids: improving the description of α/γ conformers. Biophysical journal 92, 3817–3829

42. Wang, J., Wang, W., Kollman, P. A., and Case, D. A. (2006) Automatic atom type and bond type perception in molecular mechanical calculations. Journal of molecular graphics and modelling 25, 247–260

43. Wang, J., Wolf, R. M., Caldwell, J. W., Kollman, P. A., and Case, D. A. (2004) Development and testing of a general amber force field. Journal of computational chemistry 25, 1157–1174

44. Wang, J., Wang, W., Kollman, P. A., and Case, D. A. (2001) Antechamber: an accessory software package for molecular mechanical calculations. J. Am. Chem. Soc 222, U403

45. Salomon-Ferrer, R., Götz, A. W., Poole, D., Le Grand, S., and Walker, R. C. (2013) Routine Microsecond Molecular Dynamics Simulations with AMBER on GPUs. 2. Explicit Solvent Particle Mesh Ewald. Journal of Chemical Theory and Computation 9, 3878–3888

46. Götz, A. W., Williamson, M. J., Xu, D., Poole, D., Le Grand, S., and Walker, R. C. (2012) Routine Microsecond Molecular Dynamics Simulations with AMBER on GPUs. 1. Generalized Born. Journal of Chemical Theory and Computation 8, 1542–1555

47. Ryckaert, J.-P., Ciccotti, G., and Berendsen, H. J. (1977) Numerical integration of the cartesian equations of motion of a system with constraints: molecular dynamics of n-alkanes. Journal of Computational Physics 23, 327–341

48. Roe, D. R., and Cheatham III, T. E. (2013) PTRAJ and CPPTRAJ: software for processing and analysis of molecular dynamics trajectory data. Journal of Chemical Theory and Computation 9, 3084–3095

49. Glykos, N. M. (2006) Carma: a molecular dynamics analysis program. J. Comput. Chem. 27, 1765–1768

50. Eargle, J., and Luthey-Schulten, Z. (2012) NetworkView: 3D display and analysis of protein·RNA interaction networks. Bioinformatics 28, 3000–3001

51. Sethi, A., Eargle, J., Black, A. A., and Luthey-Schulten, Z. (2009) Dynamical networks in tRNA:protein complexes. Proceedings of the National Academy of Sciences of the United States of America 106, 6620–6625

52. Newman, M. E. (2006) Modularity and community structure in networks. Proceedings of the national academy of sciences 103, 8577–8582

53. Floyd, R. W. (1962) Algorithm 97: shortest path. Communications of the ACM 5, 345

54. Team, R. C. (2019) R: A language and environment for statistical computing. R Foundation for Statistical Computing, Vienna, Austria

55. Gu, Z., Eils, R., and Schlesner, M. (2016) Complex heatmaps reveal patterns and correlations in multidimensional genomic data. Bioinformatics 32, 2847–2849

56. Wickham, H. (2016) ggplot2: Elegant Graphics for Data Analysis, Springer-Verlag, New York

57. Kassambara, A. (2020) ggpubr: ‘ggplot2’ Based Publication Ready Plots.

58. Wickham, H., Averick, M., Bryan, J., Chang, W., D’Agostino McGowan, L., François, R., Grolemund, G., Hayes, A.,,, Henry, L., Hester, J., Kuhn, M., Lin Pedersen, T., Miller, E., Milton Bache, S., Müller, K., Ooms, J., Robinson, D., Seidel, D. P., Spinu, V., Takahashi, K., Vaughan, D., Wilke, C., Woo, K., and Yutani, H. (2019) Welcome to the Tidyverse. The Journal of Open Source Software

