## Supporting Information for "The nuclear receptor LRH-1 discriminates between ligands using distinct allosteric signaling circuits"

Distinct allosteric signaling circuits used by active and inactive ligands of the nuclear receptor LRH-1

**Table S1.** Number of strong suboptimal paths to the AF-H or Tif2 from other parts of LRH-1 or from the ligands. P values refers to a two-tailed, unpaired t-test comparing complexes bound to active *versus* inactive ligands.

| | Ligand to AF-H | | Helix 6 to AF-H | | B2 to AF-H | | H7 to AF-H | | Ligand to Tif2 | Helix 6 to Tif2 | B2 to Tif2 | Helix 7 to Tif2 |
|---|---|---|---|---|---|---|---|---|---|---|---|---|
| | No coregulator | Tif2 | No coregulator | Tif2 | No coregulator | Tif2 | No coregulator | Tif2 | Tif2 | Tif2 | Tif2 | Tif2 |
| **APO** | n/a | n/a | 1230 | 2673 | 961 | 2283 | 1113 | 2251 | n/a | 5015 | 2802 | 9909 |
| **INACTIVE** | | | | | | | | | | | | |
| 1N | 22 | 125 | 118 | 269 | 55 | 1060 | 174 | 170 | 101 | 4778 | 1590 | 5608 |
| 1X | 144 | 80 | 1012 | 673 | 511 | 268 | 124 | 1107 | 48 | 1044 | 1133 | 10530 |
| 3N | 32 | 70 | 389 | 121 | 91 | 380 | 385 | 569 | 137 | 282 | 1314 | 1195 |
| 3X | 53 | 31 | 307 | 398 | 57 | 289 | 138 | 208 | 9 | 253 | 60 | 226 |
| 4X | 78 | 5 | 508 | 109 | 58 | 90 | 2511 | 307 | 21 | 139 | 226 | 1053 |
| 7X | 74 | 84 | 1992 | 166 | 96 | 709 | 1818 | 1251 | 40 | 582 | 908 | 3246 |
| 2X | 82 | 46 | 2069 | 72 | 257 | 30 | 951 | 238 | 28 | 231 | 26 | 2449 |
| Endo | 160 | 93 | 131 | 578 | 81 | 320 | 252 | 404 | 86 | 609 | 669 | 5938 |
| S1 | 101 | 10 | 1045 | 591 | 184 | 624 | 1854 | 562 | 95 | 2103 | 332 | 2451 |
| S2N | 97 | 154 | 368 | 426 | 211 | 222 | 300 | 684 | 216 | 316 | 516 | 3487 |
| S2X | 321 | 41 | 8691 | 1574 | 1148 | 274 | 635 | 150 | 23 | 327 | 176 | 2295 |
| S4X | 42 | 37 | 575 | 623 | 71 | 55 | 1012 | 427 | 12 | 39 | 14 | 29 |
| Mean | 100.5 | 64.7 | 1433.8 | 466.7 | 235.0 | 360.1 | 846.2 | 506.4 | 68.0 | 891.9 | 580.3 | 3208.9 |
| SEM | 22.4 | 12.5 | 657.4 | 113.7 | 87.4 | 83.3 | 222.9 | 99.0 | 17.2 | 372.0 | 149.8 | 819.0 |
| **ACTIVE** | | | | | | | | | | | | |
| 7N | 21 | 29 | 30 | 27 | 126 | 114 | 92 | 67 | 120 | 105 | 51 | 195 |
| 18A | 44 | 160 | 507 | 887 | 50 | 228 | 487 | 522 | 363 | 2331 | 670 | 7816 |
| 5N | 48 | 89 | 103 | 330 | 227 | 247 | 447 | 158 | 89 | 745 | 342 | 1427 |
| 5X | 46 | 66 | 85 | 546 | 152 | 353 | 184 | 229 | 60 | 169 | 886 | 212 |
| 6N | 114 | 18 | 1304 | 91 | 121 | 18 | 331 | 285 | 40 | 131 | 8 | 304 |
| 6X | 10 | 29 | 39 | 93 | 189 | 435 | 50 | 266 | 84 | 233 | 635 | 368 |
| 2N | 7 | 15 | 227 | 194 | 58 | 203 | 543 | 245 | 98 | 417 | 644 | 558 |
| RJW | 24 | 31 | 169 | 546 | 100 | 229 | 68 | 465 | 21 | 531 | 106 | 3668 |
| S3N | 35 | 29 | 560 | 1749 | 506 | 90 | 106 | 1777 | 49 | 2040 | 326 | 3340 |
| S3X | 108 | 25 | 2298 | 422 | 115 | 432 | 1104 | 215 | 31 | 135 | 220 | 922 |
| S4N | 68 | 28 | 571 | 3199 | 84 | 373 | 105 | 901 | 44 | 76 | 1594 | 4355 |
| 4N | 18 | 24 | 470 | 421 | 78 | 84 | 185 | 193 | 40 | 3227 | 1021 | 2096 |
| Mean | 45.3 | 45.3 | 530.3 | 708.8 | 150.5 | 233.8 | 308.5 | 443.6 | 86.6 | 845.0 | 541.9 | 2105.1 |
| SEM | 9.8 | 11.6 | 183.1 | 252.3 | 34.1 | 39.0 | 84.3 | 131.1 | 25.5 | 295.6 | 129.0 | 642.2 |
| p-value | 0.04 | 0.29 | 0.22 | 0.41 | 0.40 | 0.20 | 0.04 | 0.72 | 0.57 | 0.93 | 0.85 | 0.32 |

Figure S1

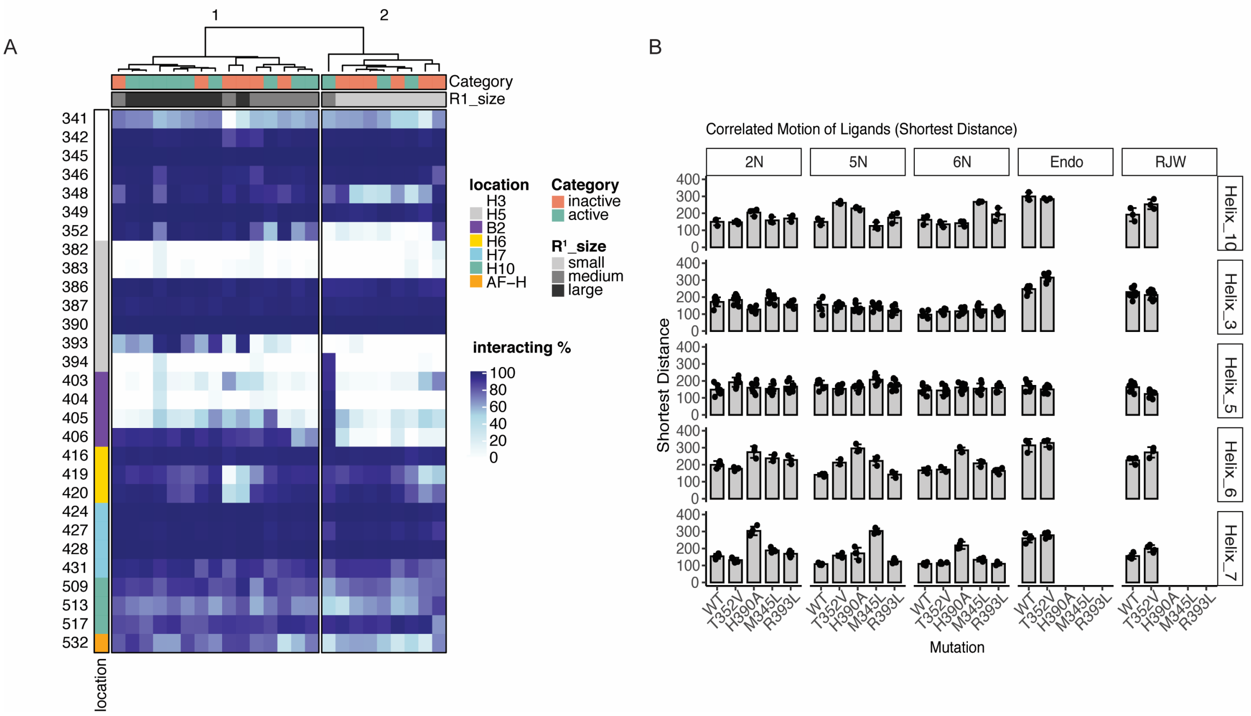

**Figure S1. Interactions made by LRH-1 ligands.** A. Heatmap showing the stability of interactions made by each ligand. Each column is a ligand, and each row is an amino acid (sequence numbers and regions of LRH-1 are indicated in the left annotation). Stability refers to the percentage of time each interaction was maintained over the course of the simulation. B. Shortest distance values for selected ligands when bound to wild-type (WT) or mutated LRH-1.

Figure S2

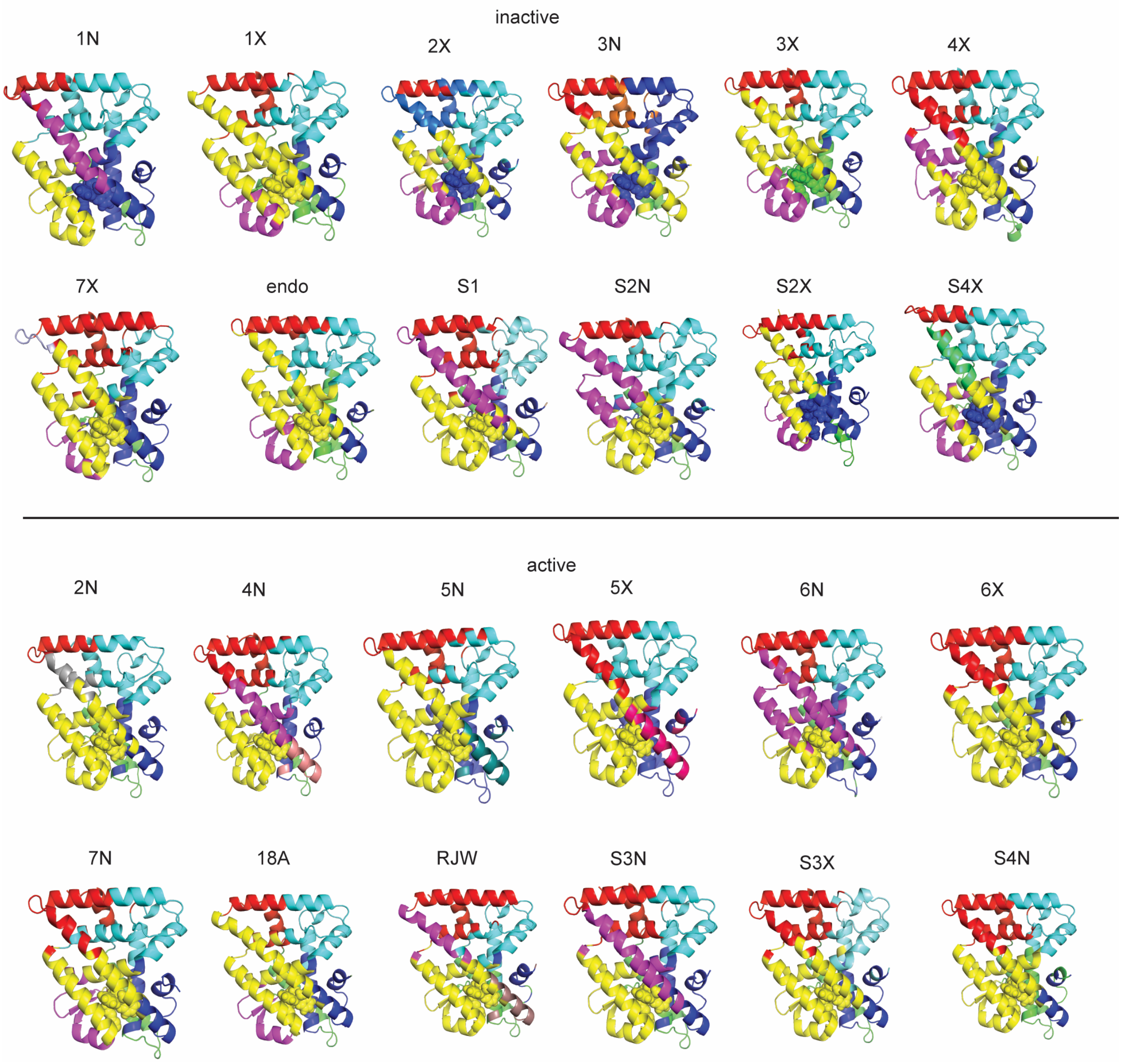

**Figure S2. Community analysis.** LRH-1 models with each ligand, colored by community. Ligands are shown as spheres.

Figure S3

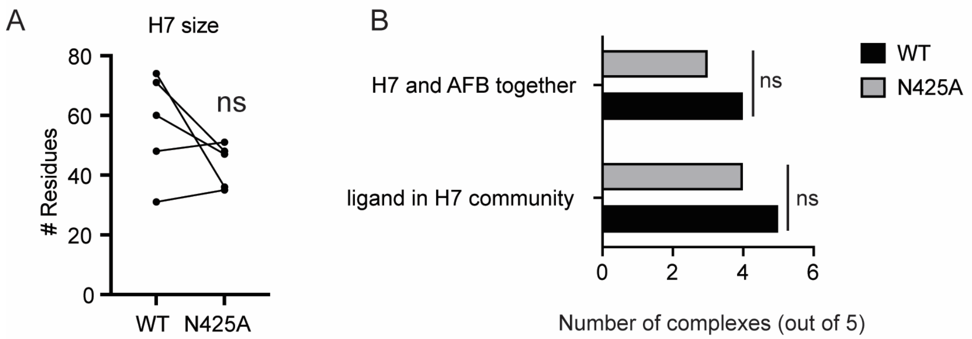

**Figure S3. Mutation of residue N425 does not significantly change helix 7 communities for a subset of ligands.** A. Size of helix7 (H7)-containing communities for wild-type (WT) or N425A LRH-1 bound to a subset of active ligands ($n = 5$). Differences were not statistically significantly different between groups ($p = 0.17$ by two-tailed, paired Student's t-test). B. The N425A mutation does not split H7 and AFB into separate communities and does not change the ligand participation in the H7 community for a subset of 5 active ligands ($p > 0.99$ for both comparisons by Fisher's exact test).
